# A guaranteed-convergence algorithm for coupled leaf photosynthesis–transpiration–stomatal conductance models

**DOI:** 10.64898/2026.06.24.734164

**Authors:** Yuji Masutomi, Kazuhiko Kobayashi

## Abstract

The photosynthesis–transpiration–stomatal conductance (*A*_*n*_–*E*–*g*_*s*_) model framework is widely used for estimating photosynthesis, transpiration, and stomatal conductance in plants. The model equations are solved by numerical iteration, and the converged model values are deemed the solution. However, there has been no general guarantee that the iterative procedure converges to a solution or that the procedure leads to convergence. Building on the recent proof of the existence of a unique set of solutions, we herewith propose a numerical algorithm that is guaranteed to converge to the solution for the *A*_*n*_–*E*–*g*_*s*_ model framework. We first analytically prove that the proposed algorithm necessarily converges to a solution. We then demonstrate the convergence across contrasting combinations of leaf temperature, relative humidity, light, atmospheric CO_2_, and wind speed. We further demonstrate rapid convergence with the algorithm: no more than ca. 10 iterations for approximately 10^−3^ *μ*mol CO_2_ m^−2^ s^−1^ precision in net photosynthesis and no more than ca. 20 iterations for 10^−7^ *μ*mol CO_2_ m^−2^ s^−1^ precision. By guaranteeing convergence to the solution, this algorithm eliminates concerns about nonconvergence in leaf gas-exchange calculations and is expected to serve as a robust foundation for a range of studies from leaf-level gas exchange to global-scale carbon and water cycle dynamics.

## 1 Introduction

Photosynthesis and transpiration by terrestrial plants play central roles in the global carbon and water cycles. Photosynthesis determines the uptake of atmospheric CO_2_ (Friedlingstein et al., 2026), while transpiration accounts for a large fraction of water returned from land to the atmosphere (Jasechko et al., 2013; Schlesinger and Jasechko, 2014). These two fluxes are tightly linked at the leaf scale because stomata regulate both CO_2_ uptake and water loss (Jones, 2014). As a result, photosynthesis, transpiration, and stomatal conductance cannot be treated as independent quantities; they must be estimated consistently.

The *A*_*n*_–*E*–*g*_*s*_ model provides a useful framework for this purpose. This formulation combines biochemical photosynthesis (Farquhar et al., 1980), stomatal regulation (Ball et al., 1987; Leuning, 1995; Medlyn et al., 2011), and CO_2_ and H_2_O diffusion (Campbell and Norman, 1998), and thereby represents the mutual dependence among carbon gain, water loss, stomatal opening, and internal CO_2_ availability. Hence, net photosynthesis *A*_*n*_, transpiration *E*, and stomatal conductance *g*_*s*_ are solved simultaneously, as well as leaf-surface CO_2_, intercellular CO_2_, and leaf-surface vapor pressure.

The *A*_*n*_–*E*–*g*_*s*_ model was explicitly formulated by Sellers et al. (1992), based on the framework proposed by Collatz et al. (1991). Since then, it has been widely used in ecosystem, crop, hydrology, climate, and Earth-system modelling. For example, it is implemented in ecosystem and land-surface models, including the Simple Biosphere Model version 2 (SiB2) (Sellers et al., 1992), the Ecosystem Demography model (Longo et al., 2019), and the Community Land Model (CLM) (Lawrence et al., 2019). As the land component of the Community Earth System Model, CLM is also used in climate and Earth-system modelling (Lawrence et al., 2019; Danabasoglu et al., 2020). The formulation has also recently been introduced into the crop model MATCRO (Masutomi et al., 2016a,b) and, together with MATCRO, into the hydrology model Integrated Land Simulator (Nitta et al., 2020; Masutomi et al., 2026).

However, solving the *A*_*n*_–*E*–*g*_*s*_ model is not straightforward. The coupled equations can be reduced to a fifth-degree polynomial (Longo et al., 2019; Masutomi, 2023), for which no general algebraic closed-form solution is available; the model therefore has to be solved numerically in practice. In most applications, the solution is obtained by numerical iteration, and the value reached after convergence is treated as the model solution. Despite this common practice, there has been no general guarantee that such iterations converge to a solution, and it is known that they may fail to converge in practice (Baldocchi, 1994; Sun et al., 2012; Lawrence et al., 2020). In CLM5, a relatively complex algorithm is used in which an alternative fallback method is invoked when the main iteration fails to converge (Lawrence et al., 2020). For other models, it is generally unclear what algorithms are used to handle nonconvergence, which poses a major problem for the reliability of estimated photosynthesis and transpiration. To date, no algorithm whose convergence is guaranteed has been proposed for this coupled model.

The objective of this study is to develop a guaranteed-convergence numerical algorithm for the *A*_*n*_–*E*–*g*_*s*_ model as a technical advance for leaf gas-exchange calculations. We analytically prove that the proposed algorithm necessarily converges to a solution by using the existence–uniqueness theorem and two model-equivalent functions presented by Masutomi and Kobayashi (2026). We then demonstrate convergence across contrasting leaf temperature, relative humidity, light, atmospheric CO_2_, and wind speed. We further evaluate the convergence rate.

## 2 Materials and Methods

### 2.1 The *A*_*n*_–*E* –*g*_*s*_ model

The *A*_*n*_–*E*–*g*_*s*_ model explored here is the same as that used for C3 plants by Lawrence et al. (2020). In this model, the net photosynthesis rate is expressed as a function of the intercellular CO_2_ concentration *C*_*i*_ by taking the minimum of the Rubisco-limited, RuBP-regeneration-limited, and triose phosphate utilization (TPU)-limited carboxylation rates, following the Farquhar photosynthesis framework (Farquhar et al., 1980):

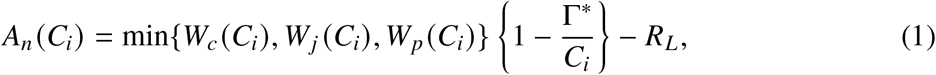

where

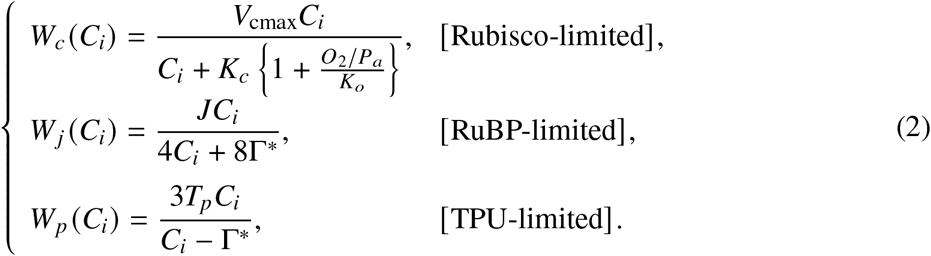

Here, *A*_*n*_ is the net photosynthesis rate, and *W*_*c*_, *W* _*j*_, and *W*_*p*_ are the Rubisco-limited, RuBP-regeneration-limited, and TPU-limited carboxylation rates, respectively. Γ^∗^ is the CO_2_ compensation point in the absence of mitochondrial respiration, *R*_*L*_ is non-photorespiratory CO_2_ release in the light, *V*_cmax_ is the maximum carboxylation rate, *J* is the electron transport rate, *T*_*p*_ is the TPU rate, *K*_*c*_ and *K*_*o*_ are Michaelis–Menten constants for CO_2_ and O_2_, *O*_2_ is the atmospheric partial pressure of oxygen, and *P*_*a*_ is atmospheric pressure. The quantities Γ^∗^, *R*_*L*_, *V*_cmax_, *K*_*c*_, and *K*_*o*_ are prescribed as functions of leaf temperature *T*_*l*_, as given in Appendix A, and *T*_*p*_ is set to *V*_cmax_/6. The electron transport rate *J* is prescribed as a function of *T*_*l*_ and absorbed photosynthetic photon flux density *Q* _*p*_, also as given in Appendix A.

In this study, we treat the three regimes separately. This is because calculating *A*_*n*_ in practice requires the regime-specific values of *W*_*c*_, *W*_*j*_, and *W*_*p*_, after which *A*_*n*_ is obtained using Eq. (1) or a smooth minimum function (Collatz et al., 1991).

Various stomatal conductance models have been proposed (Damour et al., 2010). Here, stomatal conductance for water vapor, *g*_*s*_, is represented during daytime photosynthesis by the Medlyn model (Medlyn et al., 2011), which was derived from the water-use-efficiency optimization theory of Cowan and Farquhar (1977). The Medlyn model has recently been introduced into land-surface models (Lawrence et al., 2019; Oliver et al., 2022). Under nonpositive photosynthesis, *g*_*s*_ is set to the minimum conductance. Thus, *g*_*s*_ is given by

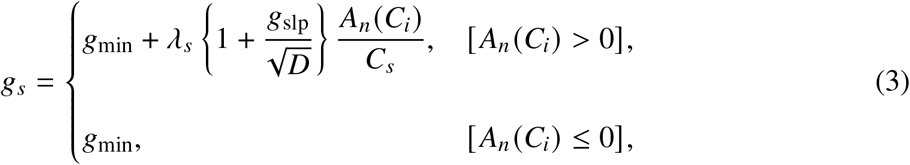

where *C*_*s*_ is the leaf-surface CO_2_ concentration, *D* is the vapor pressure deficit at the leaf surface, *g*_min_ is the minimum stomatal conductance, *g*_slp_ controls the sensitivity of conductance to photosynthesis, CO_2_, and vapor pressure deficit, and *λ*_*s*_ = 1.6 is the ratio of diffusivities of CO_2_ and water vapor through stomata. The vapor pressure deficit *D* is given by

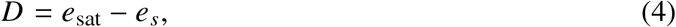

where *e*_sat_ is the saturated vapor pressure at the leaf surface and inside the leaf, and *e*_*s*_ is the vapor pressure at the leaf surface.

Following standard leaf gas-exchange mass transport relationships (Campbell and Norman, 1998), if the system is assumed to be at steady state, the CO_2_ diffusion equations connect *A*_*n*_ (*C*_*i*_), *C*_*i*_, *C*_*s*_, and the atmospheric CO_2_ concentration *C*_*a*_:

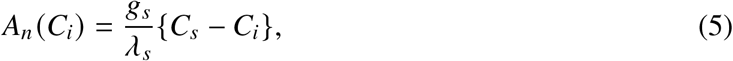

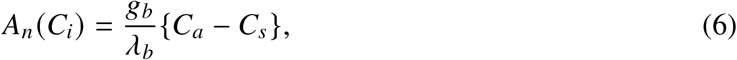

where *g*_*b*_ is the boundary-layer conductance for water vapor and *λ*_*b*_ = 1.4 is the ratio of diffusivities of CO_2_ and water vapor at the leaf surface. The boundary-layer conductance *g*_*b*_ is calculated from 2-m wind speed *u*, atmospheric pressure *P*_*a*_, and canopy height *H*, as described in Appendix B.

The corresponding water-vapor diffusion equations are

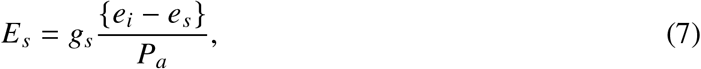

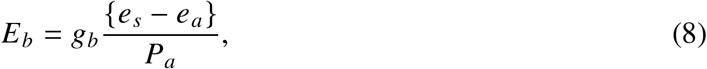

where *E*_*s*_ and *E*_*b*_ are vapor fluxes through the stomata and boundary layer, respectively, *e*_*a*_ is atmospheric vapor pressure, and *e*_*i*_ is the vapor pressure inside the leaf, which is assumed to be saturated; that is, *e*_*i*_ = *e*_sat_. The steady-state water-vapor condition is

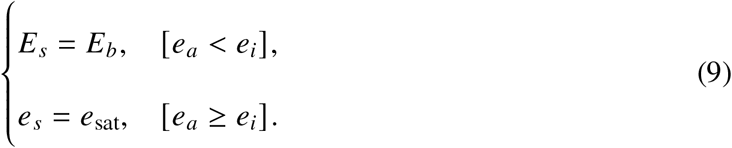

Together, Eqs. (1)–(9) define a nonlinear coupled system consisting of nine independent equation sets and nine unknown variable groups: *A*_*n*_, *W*_*c, j,p*_, *C*_*i*_, *C*_*s*_, *D, g*_*s*_, *e*_*s*_, *E*_*s*_, and *E*_*b*_. The remaining externally prescribed environmental variables and plant parameters are *C*_*a*_, *e*_*a*_, *e*_sat_, *P*_*a*_, *O*_2_, *J*, Γ^∗^, *K*_*c*_, *K*_*o*_, *R*_*L*_, *V*_cmax_, *T*_*p*_, *g*_*b*_, *g*_slp_, and *g*_min_.

### 2.2 Functions *F* (*C*_*i*_) and *G*(*C*_*i*_)

Masutomi and Kobayashi (2026) showed that the *A*_*n*_–*E*–*g*_*s*_ model can be reduced to two functions of *C*_*i*_ for stomatal conductance *g*_*s*_. These two model-equivalent functions are far more tractable, and the present paper builds the numerical algorithm on this two-function representation.

The first expression, denoted by *F* (*C*_*i*_), is obtained from the CO_2_ diffusion equations (Eqs. 5 and 6). Eliminating *C*_*s*_ from these equations gives

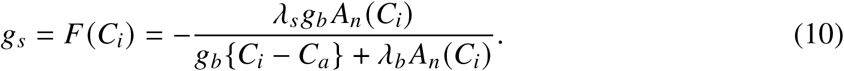

Thus, *F* (*C*_*i*_) is the stomatal conductance required by CO_2_ diffusion for a given *C*_*i*_.

For Rubisco- and RuBP-limited photosynthesis, the two limiting rates can be written in the common form

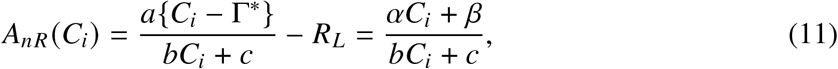

where

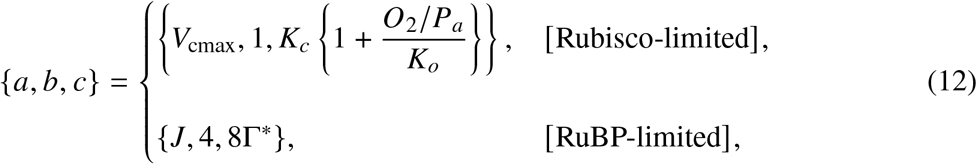

and

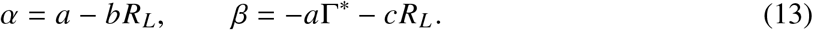

For this case,

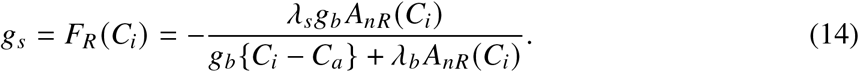

When *a* = 0, the Rubisco/RuBP-limited net photosynthesis rate reduces to

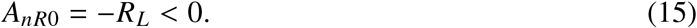

Therefore,

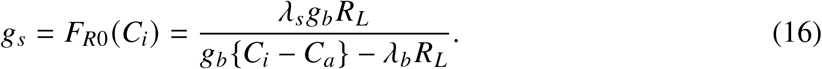

For TPU-limited photosynthesis,

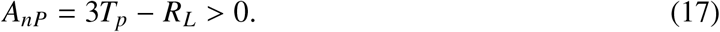

Thus,

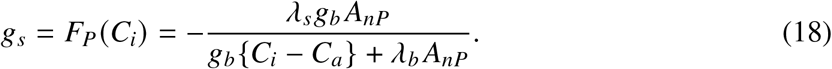

Thus, the net photosynthesis rate is summarized as

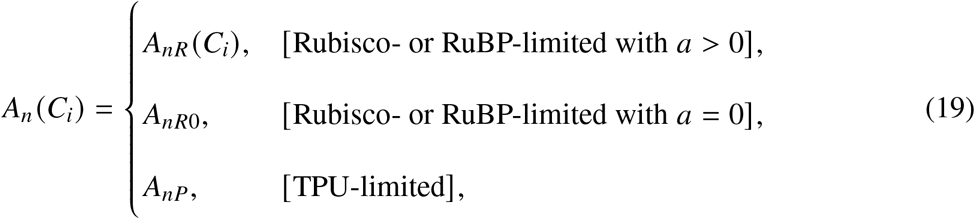

Correspondingly, the function *F* (*C*_*i*_) is given by

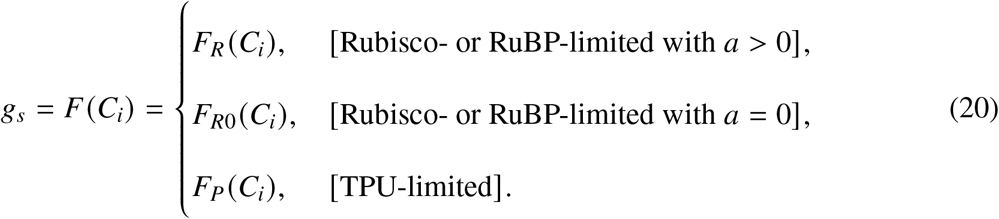

The second expression, denoted by *G*(*C*_*i*_), is obtained from the stomatal conductance model (Eq. 3), the vapor pressure deficit definition (Eq. 4), the H_2_O balance (Eqs. 7, 8, and 9), and the CO_2_ balance (Eq. 10). Eqs. (4) and (7)–(10) imply

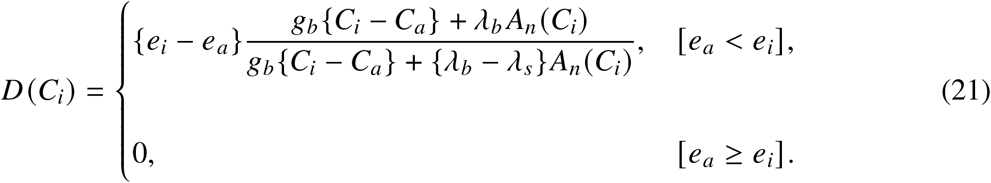

Correspondingly, *D*(*C*_*i*_) is given by

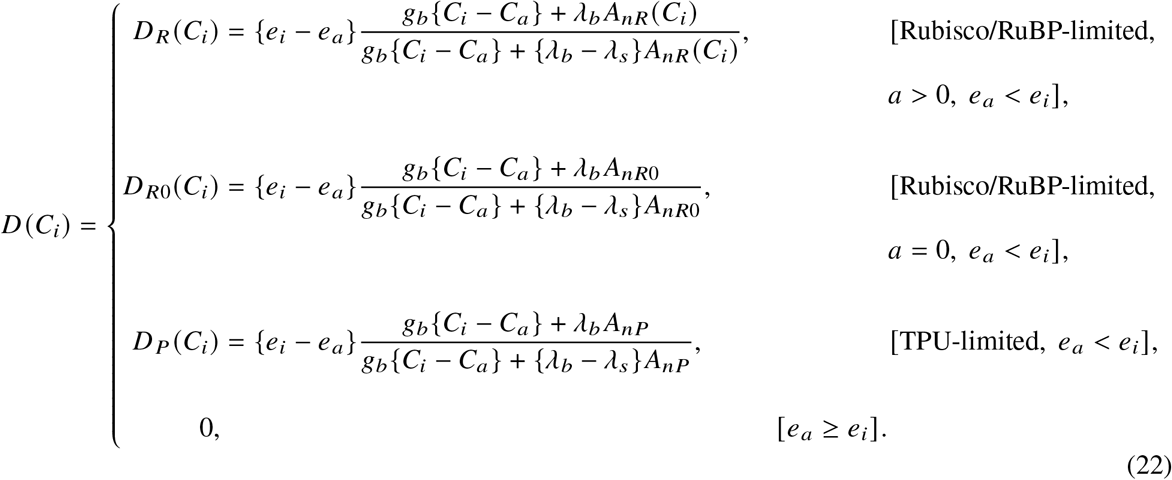

Using *D*(*C*_*i*_), the stomatal conductance model (Eq. 3) is written as

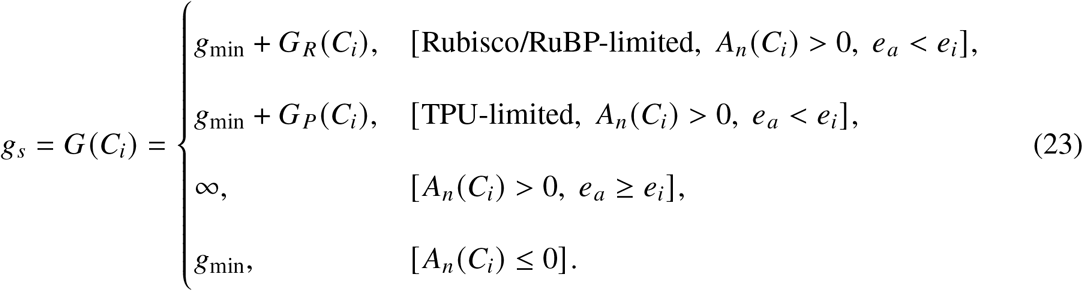

Note that *A*_*nR*0_ < 0 by Eq. (15). Therefore, the Rubisco/RuBP-limited case with *a* = 0 is included in the nonpositive-photosynthesis branch of Eq. (23). Here,

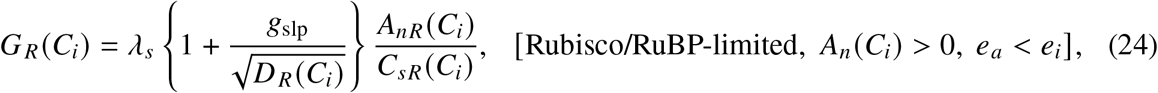

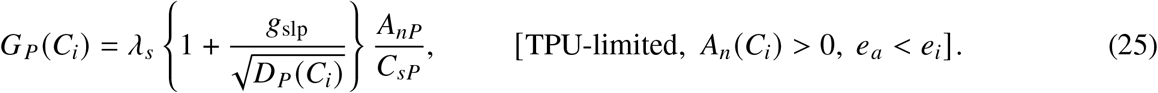

Using Eq. (6), *C*_*sR*_ (*C*_*i*_) and *C*_*sP*_ in Eqs. (24) and (25) can be expressed as follows:

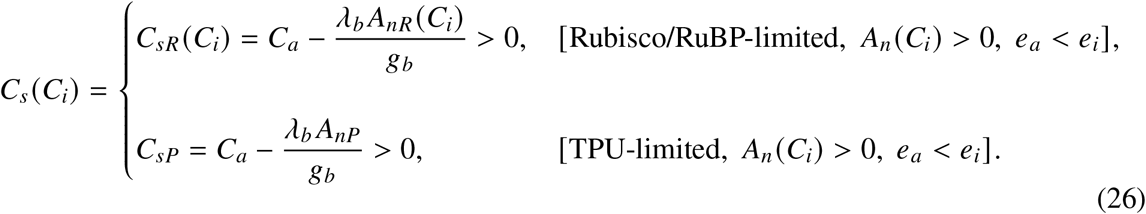

Physically, the leaf-surface CO_2_ concentration must be positive, so *C*_*s*_ (*C*_*i*_) > 0 is assumed.

### 2.3 Solution

Figure 1 gives the graphical representation of the solution of the *A*_*n*_–*E*–*g*_*s*_ model using *F* (*C*_*i*_) and *G*(*C*_*i*_). The solution is obtained at the point where these two expressions are equal, *F* (*C*_*i*_) = *G*(*C*_*i*_); the corresponding value of *C*_*i*_ defines *C*^†^, marked by the black point in Figure 1. Thus the problem of solving the multidimensional coupled model is reduced to the problem of locating the intersection of two curves in the two-dimensional *g*_*s*_–*C*_*i*_ space for each limiting condition.

**Figure 1:**
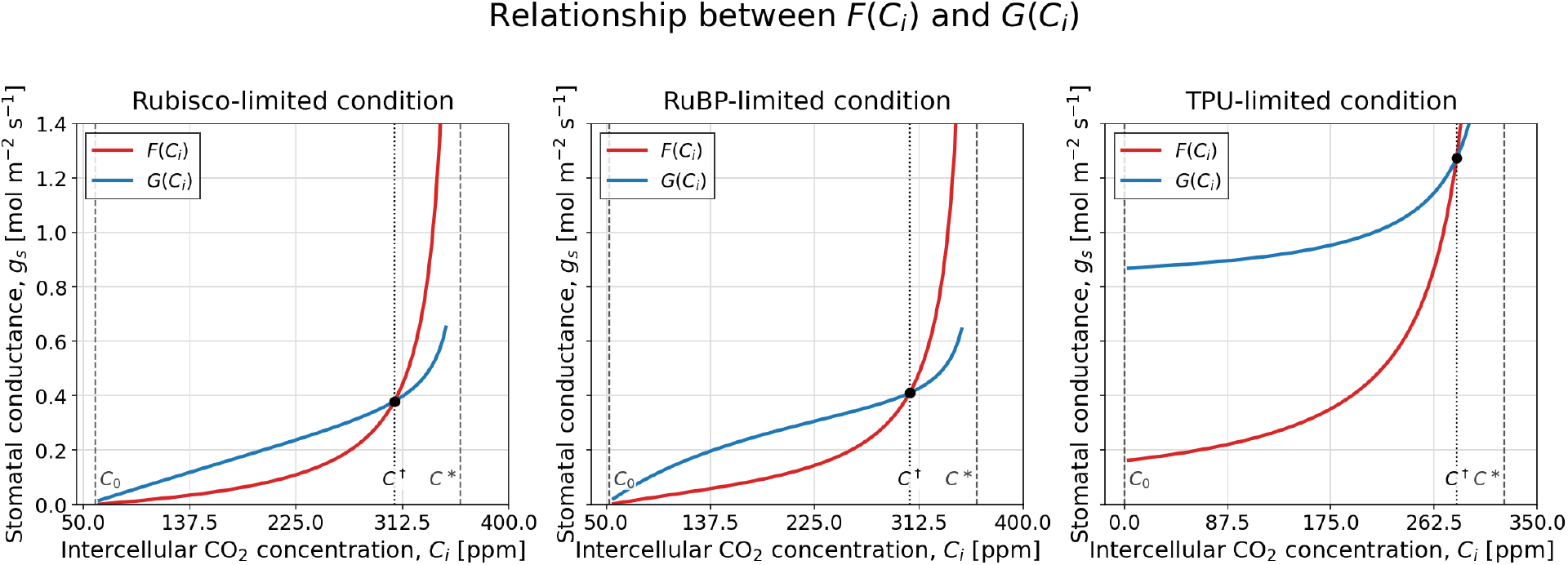
Graphical representation of the solution of the *A*_*n*_–*E*–*g*_*s*_ model using *F* (*C*_*i*_) and *G* (*C*_*i*_) for the Rubisco-limited, RuBP-limited, and TPU-limited conditions. The black point marks the solution *C*^†^ satisfying *F* (*C*^†^) = *G* (*C*^†^). Environmental inputs were *T*_*l*_ = 300 K, RH = 80%, *Q* _*p*_ = 1500 *μ*mol m^−2^ s^−1^, *u* = 1 m s^−1^, *P*_*a*_ = 101325 Pa, and *C*_*a*_ = 400 ppm.

Masutomi and Kobayashi (2026) showed that the relationships between *F* (*C*_*i*_) and *G*(*C*_*i*_) can be classified into ten geometric patterns, depending on the forms of these two functions, which vary with the associated environmental and physiological conditions. Table 1 summarizes these *F*–*G* geometric patterns and also gives the possible range of solution in *C*_*i*_; these ranges are used below to define the interval on which the numerical algorithm is applied.

**Table 1:** Geometric patterns of *F* (*C*_*i*_) and *G* (*C*_*i*_) and corresponding *C*_*i*_ ranges.

| | $F(C_i)$ | $G(C_i)$ | Limiting condition | $a$ | $A_{nR}(\pm\infty)$ | $e_a$ and $e_i$ | Range |
| --- | --- | --- | --- | --- | --- | --- | --- |
| R-I | $F_R(C_i)$ | $g_{\min} + G_R(C_i)$ | Rubisco/RuBP | $a > 0$ | $A_{nR}(\pm\infty) > 0$ | $e_a < e_i$ | $F_R^{-1}(0) < C_i < F_R^{-1}(*)^+$ |
| R-II | $F_R(C_i)$ | $g_{\min} + G_R(C_i)$ | Rubisco/RuBP | $a > 0$ | $A_{nR}(\pm\infty) > 0$ | $e_a < e_i$ | $F_R^{-1}(0) < C_i < F_R^{-1}(*)^+$ |
| R-III | $F_R(C_i)$ | $+\infty$ | Rubisco/RuBP | $a > 0$ | $A_{nR}(\pm\infty) > 0$ | $e_a \geq e_i$ | $F_R^{-1}(0) < C_i < F_R^{-1}(*)^+$ |
| R-IV | $F_R(C_i)$ | $g_{\min}$ | Rubisco/RuBP | $a > 0$ | $A_{nR}(\pm\infty) > 0$ | – | $F_R^{-1}(*)^+ < C_i < F_R^{-1}(0)$ |
| R-V | $F_R(C_i)$ | $g_{\min}$ | Rubisco/RuBP | $a > 0$ | $A_{nR}(\pm\infty) > 0$ | – | $C_i = F_R^{-1}(*)^+ = F_R^{-1}(0)$ |
| R-VI | $F_R(C_i)$ | $g_{\min}$ | Rubisco/RuBP | $a > 0$ | $A_{nR}(\pm\infty) < 0$ | – | $F_R^{-1}(*)^+ < C_i$ |
| R-VII | $F_R(C_i)$ | $g_{\min}$ | Rubisco/RuBP | $a > 0$ | $A_{nR}(\pm\infty) = 0$ | – | $F_R^{-1}(*)^+ < C_i$ |
| R-VIII | $F_{R0}(C_i)$ | $g_{\min}$ | Rubisco/RuBP | $a = 0$ | – | – | $F_{R0}^{-1}(*) < C_i$ |
| P-I | $F_P(C_i)$ | $g_{\min} + G_P(C_i)$ | TPU | – | – | $e_a < e_i$ | $0 < C_i < F_P^{-1}(*)$ |
| P-II | $F_P(C_i)$ | $+\infty$ | TPU | – | – | $e_a \geq e_i$ | $0 < C_i < F_P^{-1}(*)$ |

Among these ten patterns, the cases that require numerical solution are R-I, R-II, and P-I, because *G*(*C*_*i*_) varies with *C*_*i*_. In the other cases, *G*(*C*_*i*_) is a constant value, either *g*_min_ or the boundary value +∞, so the solution can be obtained analytically by applying the inverse *F*^−1^ (see Appendix D for the explicit expression of *F*^−1^). Because R-I and R-II have the same expressions for *F* (*C*_*i*_) and *G*(*C*_*i*_), numerical solution is essentially required for two distinct patterns, represented by R-I/II and P-I.

### 2.4 Numerical Algorithm

As described in the previous subsection, numerical solution is required for R-I/II and P-I. Here, we describe the solution algorithm for these two cases.

Figure 2 outlines how the algorithm uses these two curves to approach the solution. For example, starting from a given value 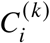, the value 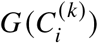 is read from the blue curve, *G*(*C*_*i*_). This conductance is then mapped back through the inverse of the red curve, *F*^−1^, to obtain the next value 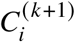. Applying the same operation repeatedly gives 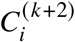 and 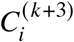, and continued application moves the sequence toward the unique intersection *C*^†^, where *F* (*C*^†^) = *G*(*C*^†^). Whether this procedure approaches *C*^†^ depends on the shapes of *F* and *G* and on the choice of the initial value; sufficient conditions for convergence are given in Section 2.5.

**Figure 2:**
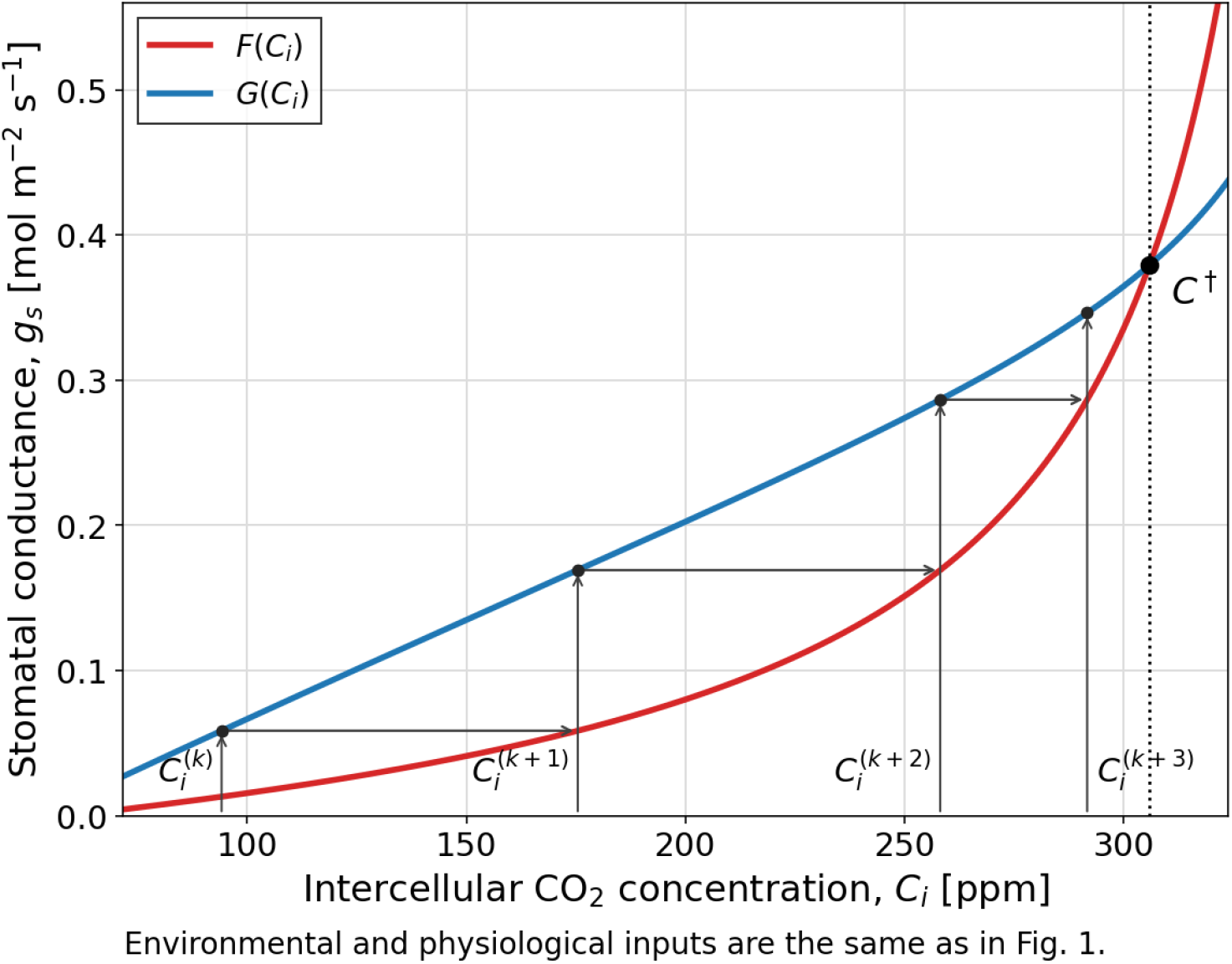
Illustration of successive steps of the proposed iteration. Starting from a current iterate 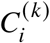, the conductance 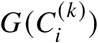 is evaluated on the blue curve, *G*(*C*_*i*_), and then mapped through *F*^−1^ to obtain 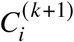 on the red curve; repeating the same operation gives 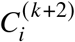 and then 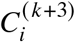. The sequence approaches the unique crossing *C*^†^, where *F* (*C*^†^) = *G*(*C*^†^).

The initial value of the iteration and the iteration interval are chosen from the possible range of solution in *C*_*i*_ shown in Table 1. For R-I and R-II, the possible range is 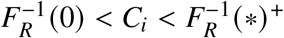, and the iteration starts from the lower endpoint 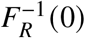. For P-I, the possible range is 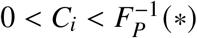, and the iteration starts from the lower endpoint 0. The concrete expressions for the zeros of *F, F*^−1^(0), and the singular points, *F*^−1^(∗), are given in Appendix C. Starting from the analytically defined endpoint avoids an arbitrary initial guess and keeps the iteration inside the specified interval of *C*_*i*_.

Using *G* and *F*^−1^, this update procedure can be written as the following one-dimensional formulation:

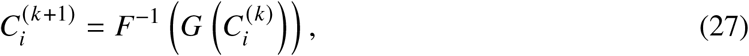

with the initial value

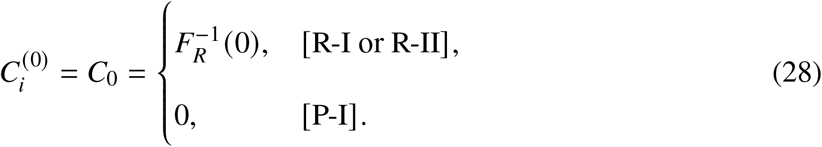

Repeated application of Eq. (27) makes the sequence approach *C*^†^, although in general the iterates do not reach *C*^†^ exactly in a finite number of steps. Therefore, in numerical calculation, the iteration is stopped when

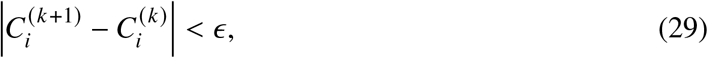

where *ϵ* is a prescribed convergence tolerance. The resulting value is then regarded as sufficiently close to *C*^†^ and is adopted as the numerical solution for *C*_*i*_. Once this value of *C*_*i*_ has been obtained, the remaining variables, including *g*_*s*_, *A*_*n*_, *C*_*s*_, *D, e*_*s*_, and *E*, are calculated by substituting it into the model equations. In this way, values satisfying all model equations are obtained.

### 2.5 Sufficient conditions for convergence

This subsection gives sufficient conditions under which the iterative procedure described in Section 2.4 makes *C*_*i*_ approach *C*^†^. Let

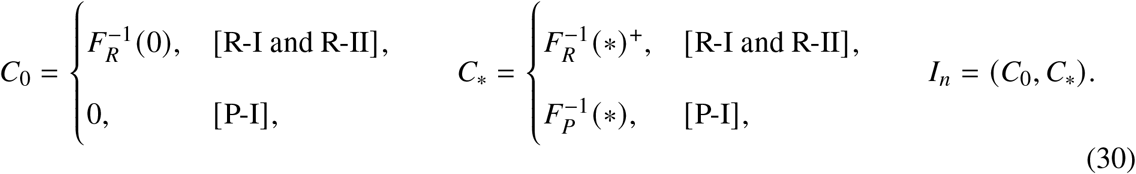

Then the following set of conditions is sufficient for the iteration 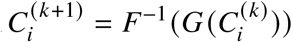, initialized at 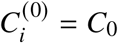, to converge monotonically to the solution.

#### Sufficient conditions for convergence

1. There is a unique crossing *C*^†^ ∈ *I*_*n*_ satisfying

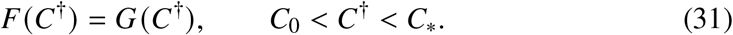
2. At the left endpoint of the possible range of solution, *F* (*C*_0_) < *G*(*C*_0_).
3. On *I*_*n*_, *F*^′^(*C*_*i*_) > 0 and *G*^′^(*C*_*i*_) > 0.

These conditions allow us to verify convergence as follows. Since *F* (*C*_0_) < *G*(*C*_0_) and the crossing is unique, continuity gives

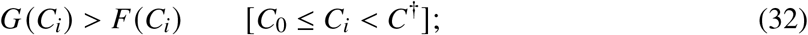

otherwise an additional crossing would occur before *C*^†^. Because *F*^′^(*C*_*i*_) > 0, Eq. (32) gives

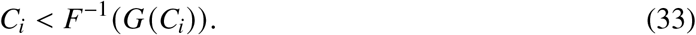

Thus each update moves to the right. Because *G*^′^(*C*_*i*_) > 0, for *C*_*i*_ < *C*^†^ we also have *G*(*C*_*i*_) < *G*(*C*^†^) = *F* (*C*^†^). Applying the increasing inverse *F*^−1^ gives

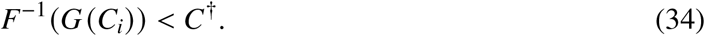

Combining Eqs. (33) and (34) yields

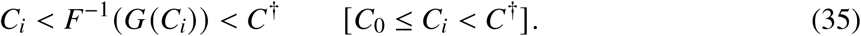

Therefore the iterates remain below the unique crossing, increase monotonically, and converge to *C*^†^.

The first condition is satisfied by the existence–uniqueness theorem of Masutomi and Kobayashi (2026). The second condition follows from the left endpoints of the possible ranges of solution in Table 1. For R-I and R-II, 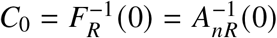, and hence *A*_*nR*_ (*C*_0_) = 0. Therefore

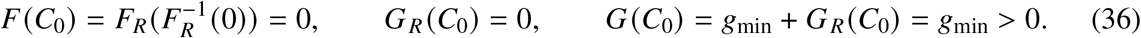

Thus *F* (*C*_0_) < *G*(*C*_0_) for R-I and R-II.

For P-I, *C*_0_ = 0. Because

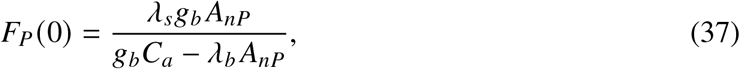

and

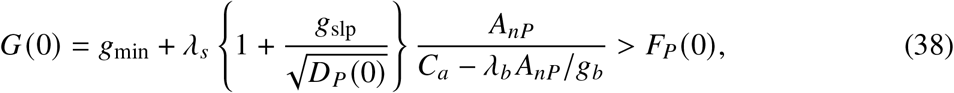

the inequality *F* (*C*_0_) < *G*(*C*_0_) holds for P-I.

The third condition has not yet been proved; the Results section proves the two monotonicity conditions *F*^′^(*C*_*i*_) > 0 and *G*^′^(*C*_*i*_) > 0.

## 3 Results

### 3.1 Proof of convergence

This subsection proves the monotonicity conditions *F*^′^(*C*_*i*_) > 0 and *G*^′^(*C*_*i*_) > 0 for R-I/II and P-I. For R-I/II, *F*_*R*_ (*C*_*i*_) is given by

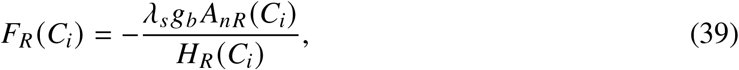

where *H*_*R*_ (*C*_*i*_) is defined as

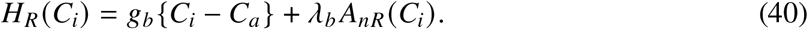

The derivative of *F*_*R*_ is

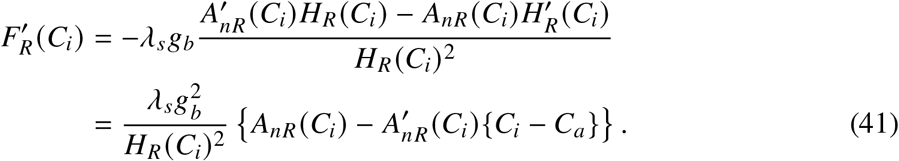

Since 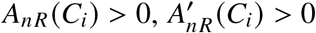, and *C*_*i*_ < *C*_*a*_ on *I*_*n*_ for R-I/II, we have

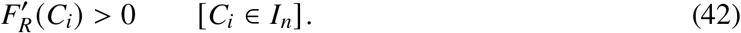

For R-I/II, *G*_*R*_ (*C*_*i*_) is given by

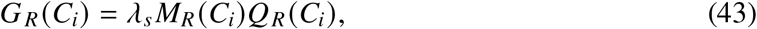

where *Q*_*R*_ (*C*_*i*_) and *M*_*R*_ (*C*_*i*_) are defined as

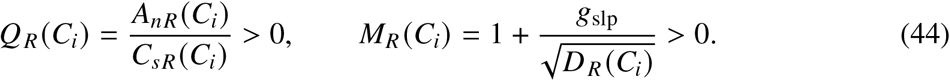

Here, Eq. (26) has been used to show *Q*_*R*_ (*C*_*i*_) > 0. The quantity *D*_*R*_ (*C*_*i*_) is given by

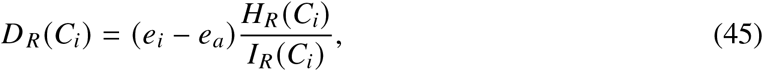

where *I*_*R*_ (*C*_*i*_) is defined as

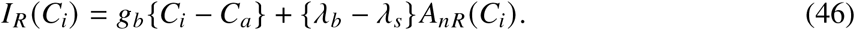

The derivative of *G*_*R*_ is

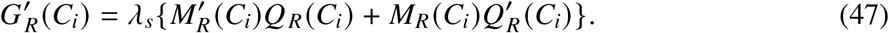

Differentiating *M*_*R*_ (*C*_*i*_) gives

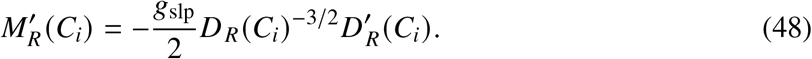

where 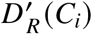 is given by

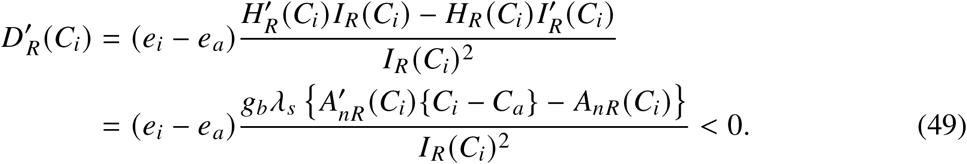

Therefore, from Eqs. (48) and (49),

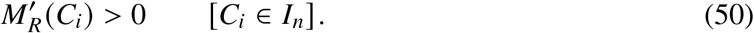

Similarly, differentiating *Q*_*R*_ (*C*_*i*_) gives

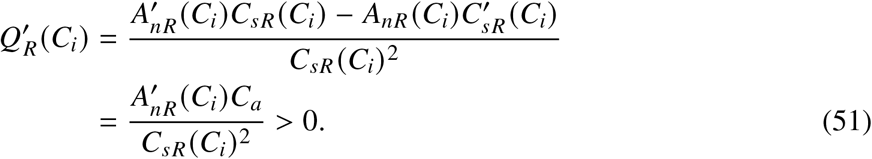

Therefore, from Eqs. (47), (44), (50), and (51),

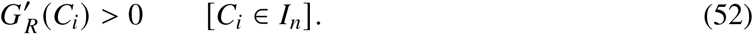

For P-I, the function *F*_*P*_ (*C*_*i*_) is given by

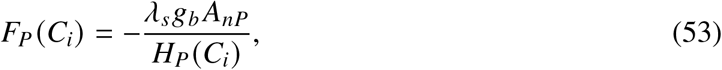

where *H*_*P*_ (*C*_*i*_) is defined as

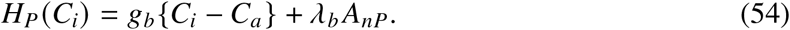

The derivative of *F*_*P*_ is

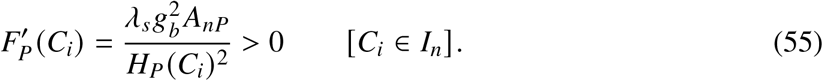

For P-I, *G*_*P*_ (*C*_*i*_) is given by

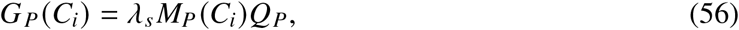

where *Q*_*P*_ and *M*_*P*_ (*C*_*i*_) are defined as

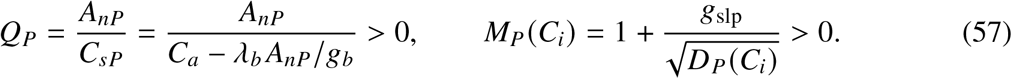

Here, Eq. (26) has been used to show *Q*_*P*_ > 0. The quantity *D*_*P*_ (*C*_*i*_) is given by

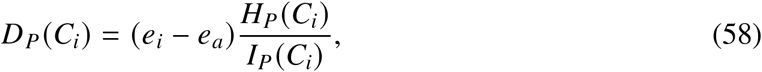

where *I*_*P*_ (*C*_*i*_) is defined as

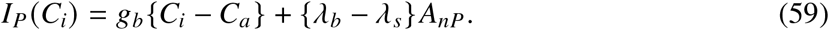

The derivative of *G*_*P*_ is

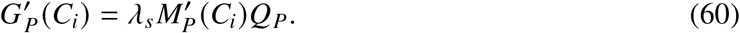

Differentiating *M*_*P*_ (*C*_*i*_) gives

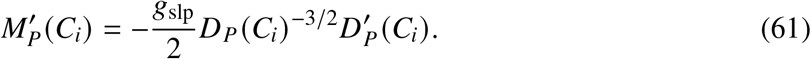

where 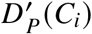 is given by

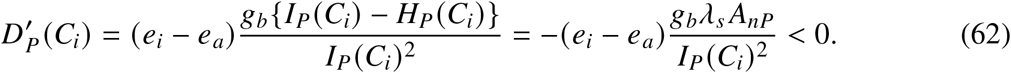

Therefore, from Eqs. (61) and (62),

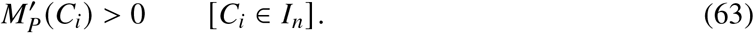

Therefore, from Eqs. (60), (57), and (63),

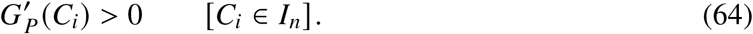

This completes the monotonicity proof required by the convergence conditions in Section 2.5.

### 3.2 Iteration behavior under contrasting environmental conditions

We next examined the numerical behavior of the proposed iteration under contrasting environmental conditions. This analysis had two purposes: first, to confirm that the sequence generated by Eq. (27) converged to the condition-specific solution in each tested case, and second, to quantify how rapidly the convergence criterion was satisfied. Unless otherwise stated, the iteration was stopped when 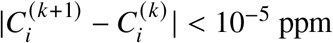.

Figures 3–7 show iteration trajectories for the Rubisco-limited, RuBP-limited, and TPU-limited conditions when one environmental input was varied at a time: leaf temperature *T*_*l*_, relative humidity *RH*, absorbed photosynthetic photon flux density *Q* _*p*_, atmospheric CO_2_ concentration *C*_*a*_, and wind speed *u*. Unless otherwise varied in each figure, the environmental inputs were fixed at the same values as in Fig. 1, and the remaining parameter values were those given in Appendices A and B.

**Figure 3:**
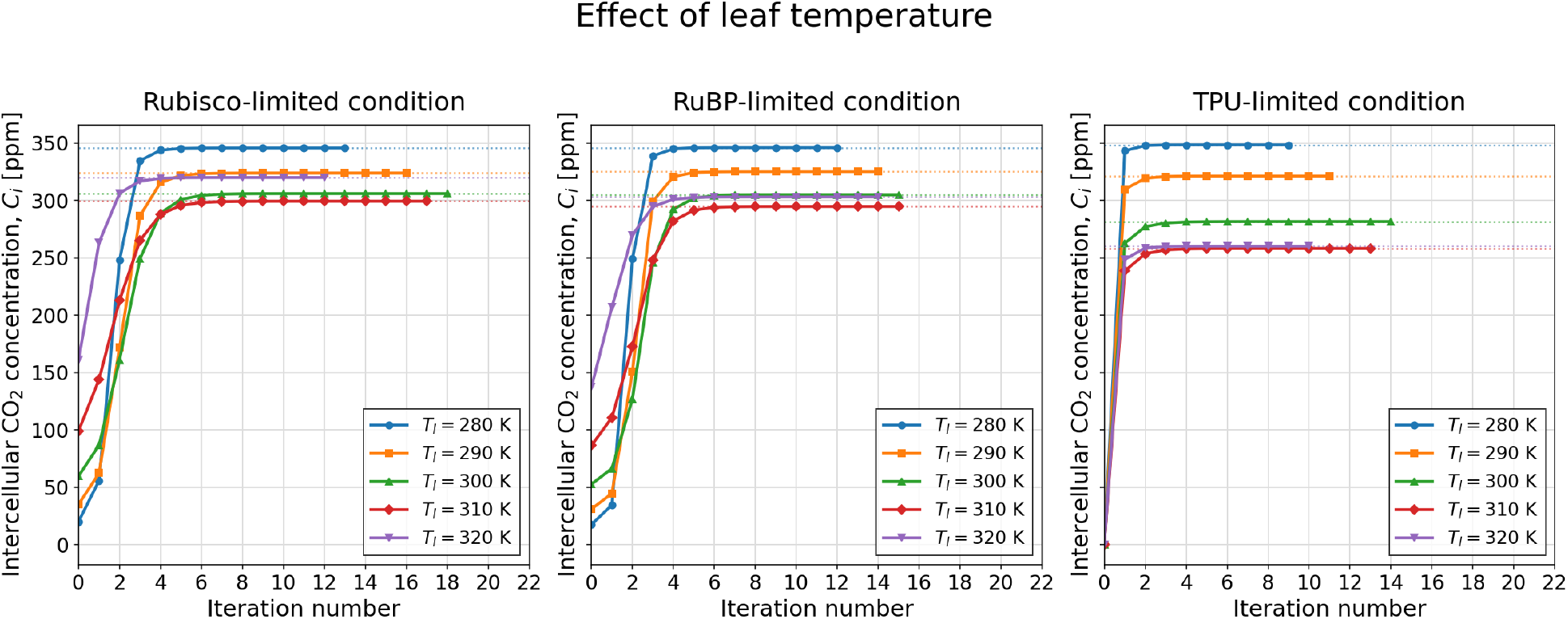
Iteration trajectories of *C*_*i*_ for the Rubisco-limited, RuBP-limited, and TPU-limited conditions when leaf temperature *T*_*l*_ is varied. Variables other than *T*_*l*_ are fixed at the Fig. 1 values.

**Figure 4:**
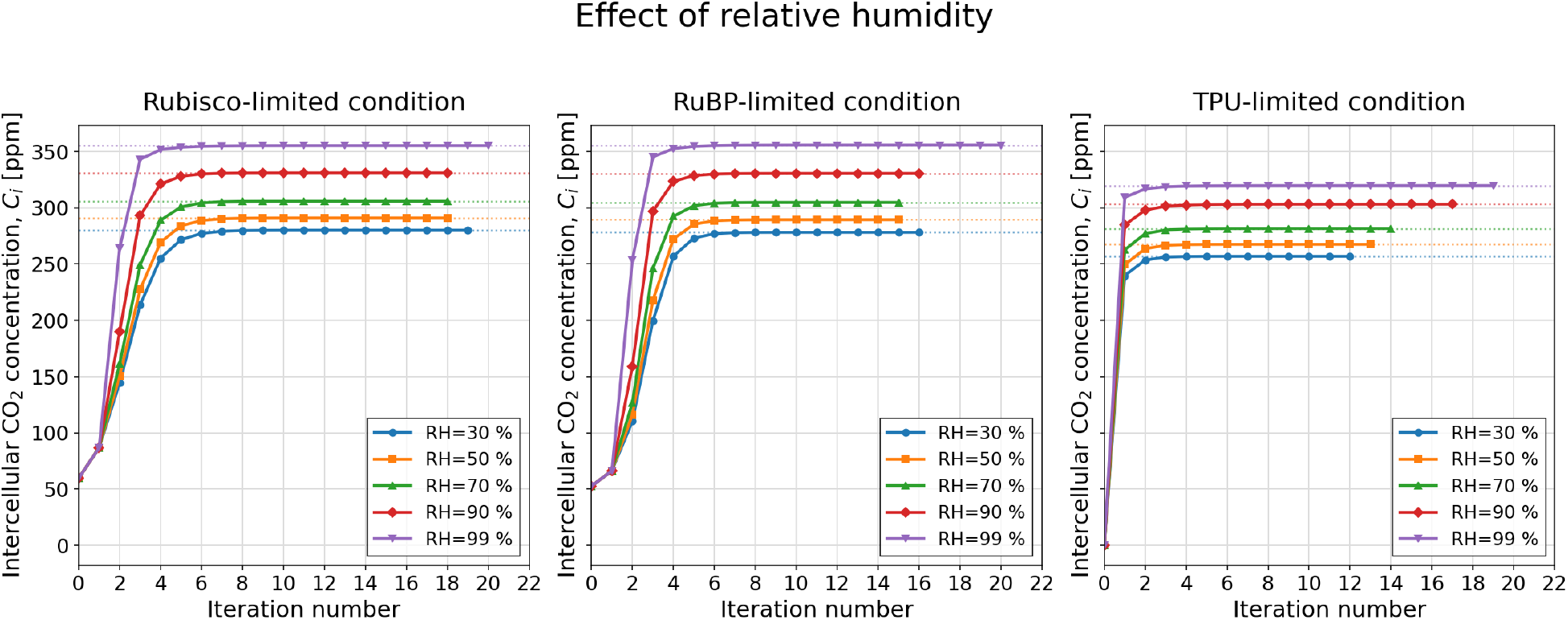
Iteration trajectories of *C*_*i*_ for the Rubisco-limited, RuBP-limited, and TPU-limited conditions when relative humidity *RH* is varied. Variables other than *RH* are fixed at the Fig. 1 values.

**Figure 5:**
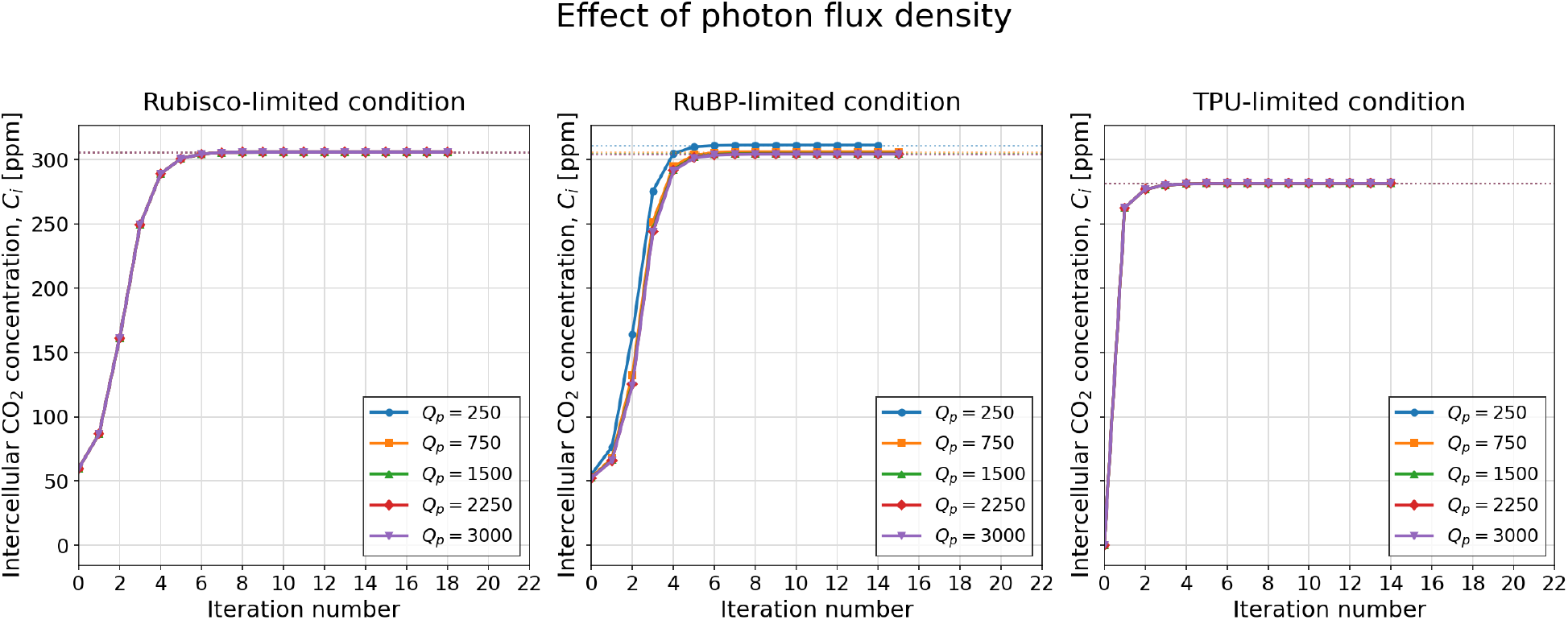
Iteration trajectories of *C*_*i*_ for the Rubisco-limited, RuBP-limited, and TPU-limited conditions when absorbed photosynthetic photon flux density *Q* _*p*_ is varied. Variables other than *Q* _*p*_ are fixed at the Fig. 1 values.

**Figure 6:**
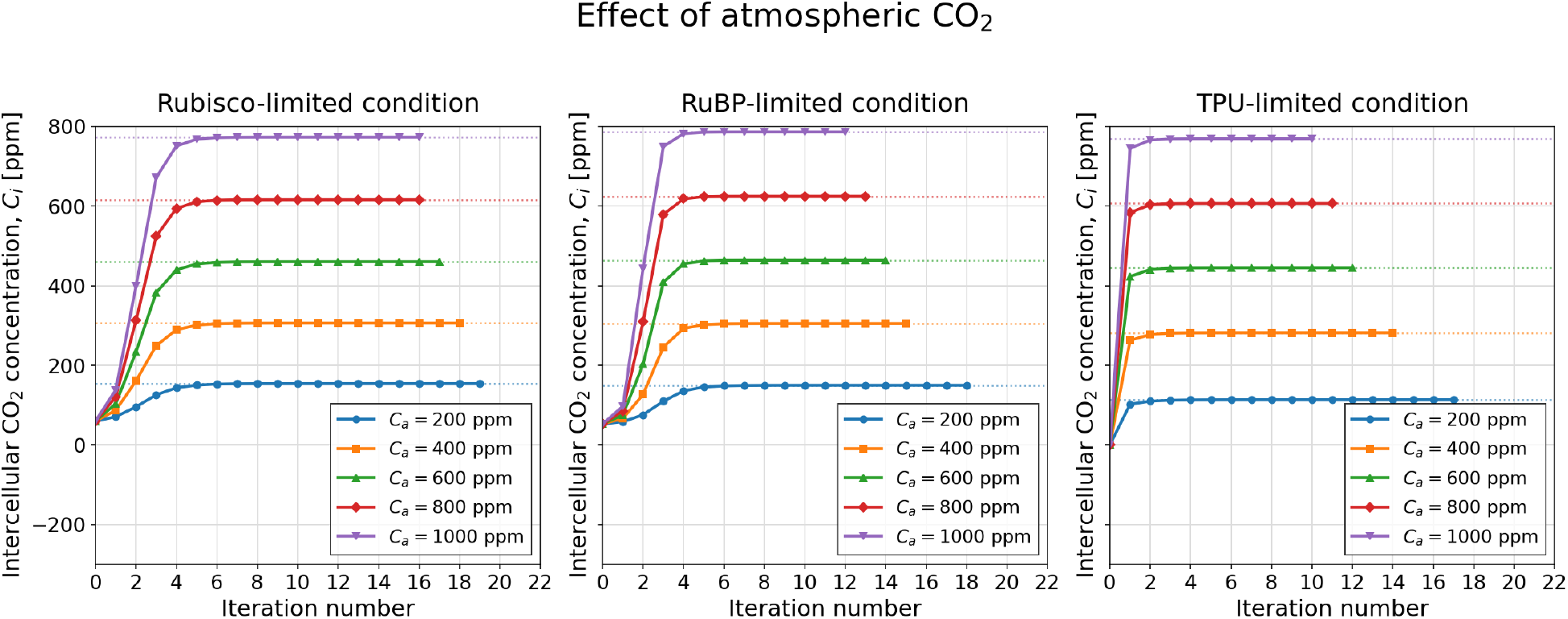
Iteration trajectories of *C*_*i*_ for the Rubisco-limited, RuBP-limited, and TPU-limited conditions when atmospheric CO_2_ concentration *C*_*a*_ is varied. Variables other than *C*_*a*_ are fixed at the Fig. 1 values.

**Figure 7:**
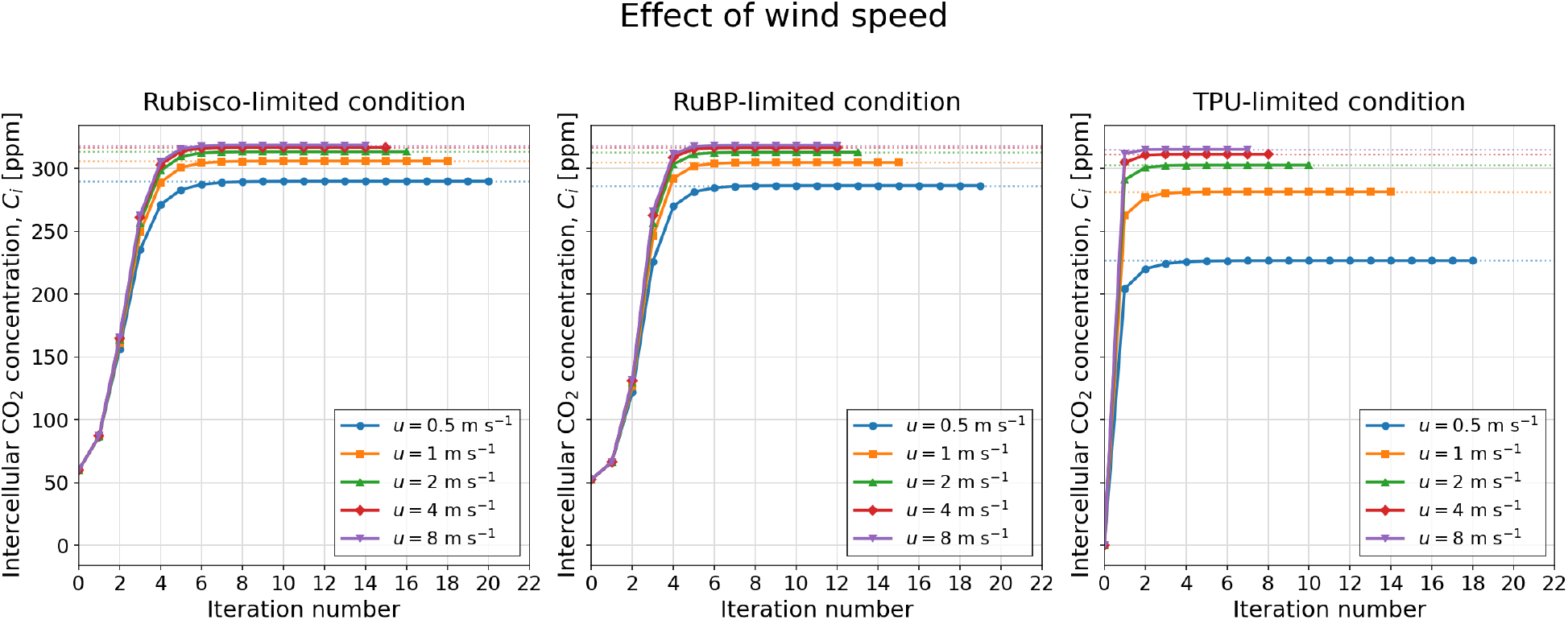
Iteration trajectories of *C*_*i*_ for the Rubisco-limited, RuBP-limited, and TPU-limited conditions when wind speed *u* is varied. Variables other than wind speed are fixed at the Fig. 1 values.

The trajectories in Figs. 3–7 first confirm that convergence was reached in all tested cases across the Rubisco-limited, RuBP-limited, and TPU-limited conditions. These trajectories also show several common features. First, convergence was monotonic in all cases shown: starting from the lower endpoint, *C*_*i*_ increased rapidly during the first few iterations and then approached a stable plateau without oscillation or overshooting, as theoretically predicted by Eq. (35). Second, although the limiting conditions differed in their converged values and in the number of iterations required, their convergence trajectories were qualitatively similar. Most of the change occurred within approximately the first four to six iterations, after which the updates became very small.

The effect of the convergence threshold itself was evaluated for the one-variable sensitivity tests separately for the Rubisco-limited, RuBP-limited, and TPU-limited conditions. The convergence threshold for 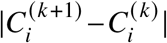 was varied as 10^−1^, 10^−2^, 10^−3^, 10^−4^, and 10^−5^ ppm. Figure 8 shows that the number of iterations required for convergence increased as the threshold became stricter across the tested environmental variables and the three limiting conditions; the corresponding threshold-specific panels are provided in Appendix Figs. A1–A5. Even so, convergence remained rapid: a precision of 0.1 ppm required no more than about 10 iterations, and even the strictest precision tested, 10^−5^ ppm, required no more than about 20 iterations in these examples. A notable feature was that the number of iterations increased approximately linearly with the number of decimal orders of precision required up to the 10^−5^ ppm threshold, indicating predictable improvement in numerical accuracy with additional iterations.

**Figure 8:**
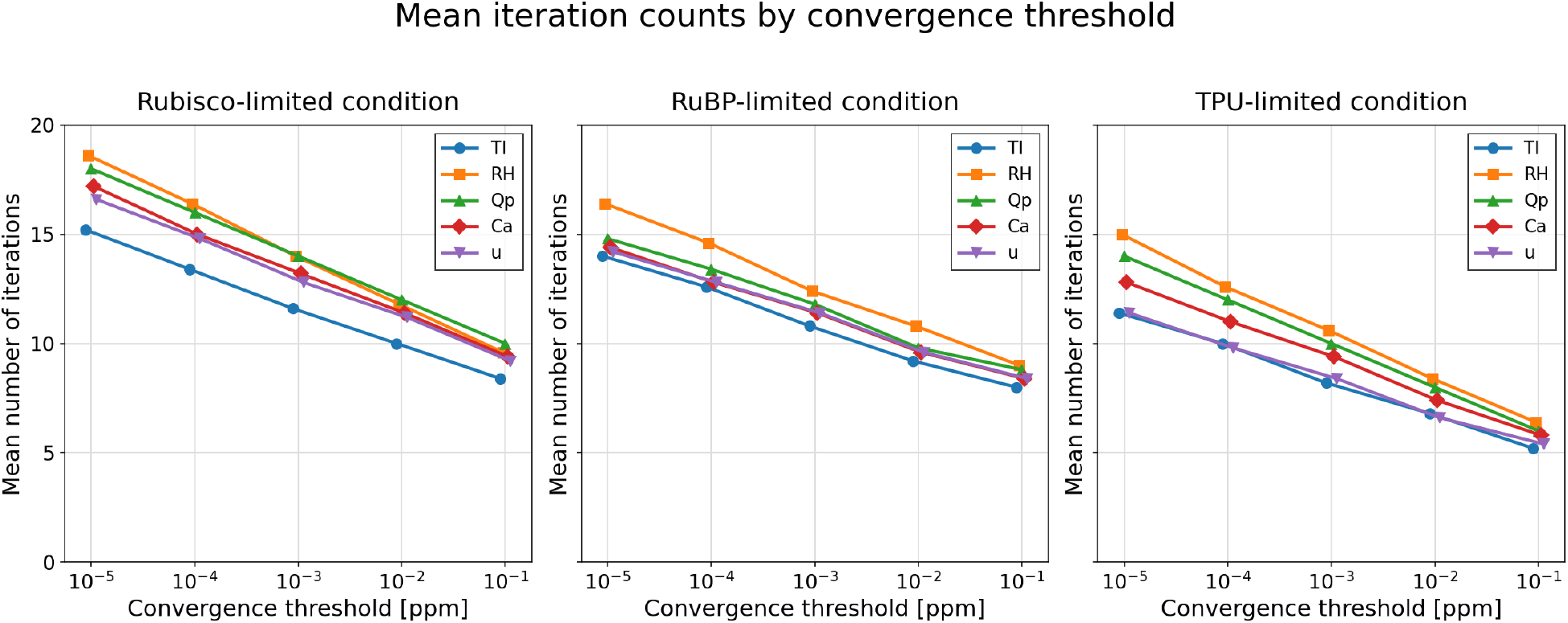
Combined summary of the iteration counts shown separately in Appendix Figs. A1–A5 for the Rubisco-limited, RuBP-limited, and TPU-limited conditions. The horizontal axis is log_10_ of the convergence threshold applied to 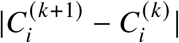, and the vertical axis is the mean number of iterations required for convergence, calculated separately for each environmental variable. Marker shape denotes the environmental variable varied in the one-variable sensitivity test. Each point shows the mean iteration count across the tested values of that variable under the corresponding convergence threshold.

We also evaluated the convergence of net photosynthesis *A*_*n*_ at the stopping point of *C*_*i*_. For each one-variable sensitivity test and convergence threshold, we calculated the final-iteration change in *A*_*n*_ for the Rubisco-limited and RuBP-limited conditions. The TPU-limited condition is omitted because *A*_*n*_ = 3*T*_*p*_ − *R*_*L*_ is independent of *C*_*i*_ within that limiting regime. Figure 9 shows that the final-iteration change in *A*_*n*_ decreased systematically as the *C*_*i*_ convergence threshold became stricter. Specifically, it was approximately 10^−3^ *μ*mol CO_2_ m^−2^ s^−1^ for the 10^−1^ ppm threshold and approximately 10^−7^ *μ*mol CO_2_ m^−2^ s^−1^ for the 10^−5^ ppm threshold. These thresholds correspond to no more than about 10 and 20 iterations, respectively, in Fig. 8.

**Figure 9:**
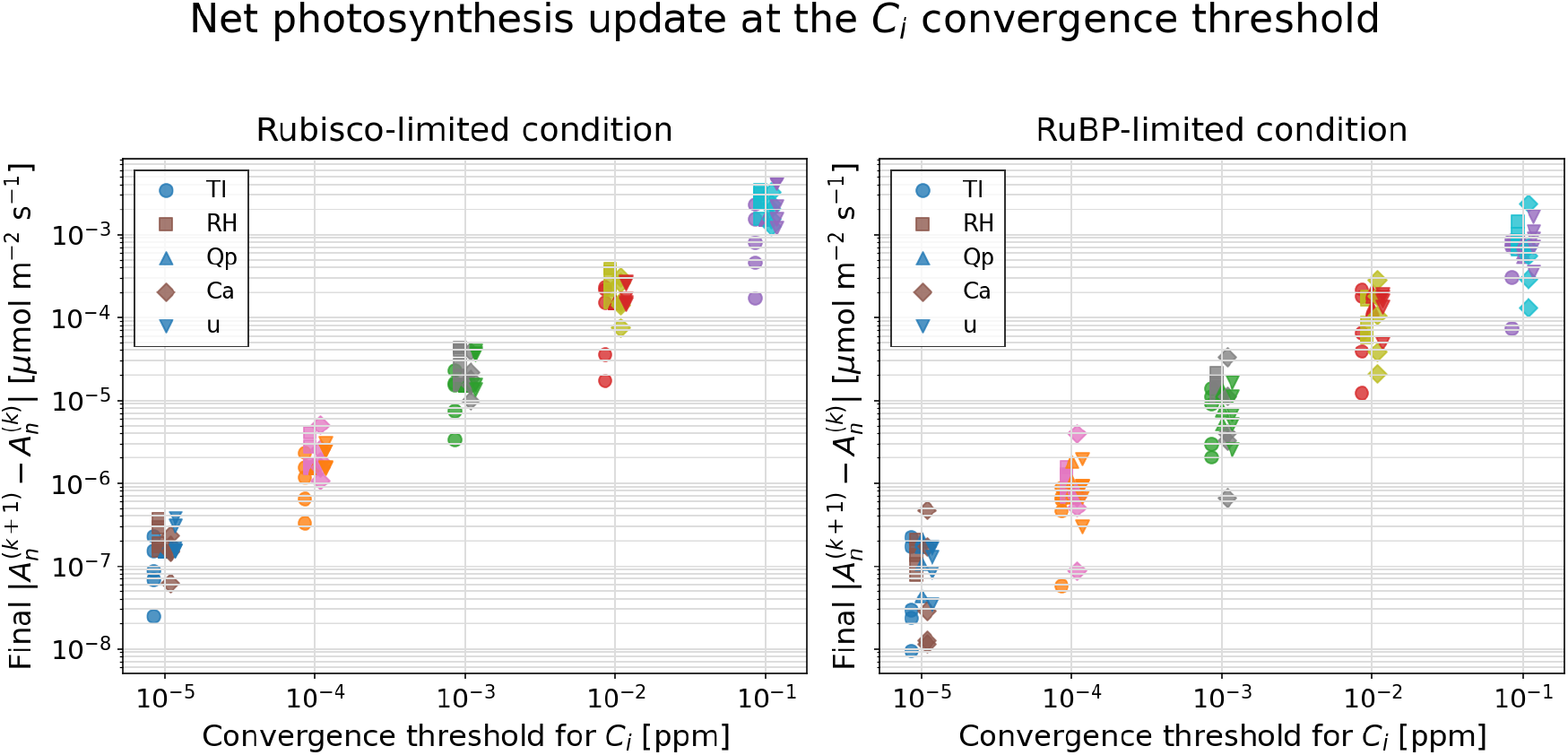
Relationship between the convergence threshold for *C*_*i*_ and the final-iteration change in *A*_*n*_ for the Rubisco-limited and RuBP-limited conditions. Each point represents one environmental value from the one-variable sensitivity tests. Marker shape denotes the environmental variable varied in the sensitivity test, and slight horizontal offsets are used for visibility.

## 4 Discussion

The main technical advance of this study is the development of a numerical algorithm whose convergence is guaranteed when solving the *A*_*n*_–*E*–*g*_*s*_ model. Since the influential work of Collatz et al. (1991) and Sellers et al. (1992), many studies and models have used numerical iterative schemes to solve the *A*_*n*_–*E*–*g*_*s*_ model. However, nonconvergence of such coupled photosynthesis– stomatal conductance calculations has been reported previously (Baldocchi, 1994; Sun et al., 2012; Lawrence et al., 2020), and when an iteration does not converge, the correct solution cannot be obtained reliably. In such cases, it is unclear how nonconvergence is handled in many models, but in CLM5, for example, a backup solver is used when the main iteration fails to converge (Lawrence et al., 2020); such a strategy is practical, but it makes the solver hybrid and more complex. Furthermore, convergence of this hybrid approach itself has not been proved. To our knowledge, this is the first algorithm for this model whose convergence is mathematically guaranteed. This property eliminates concerns about nonconvergence and provides robust and reliable leaf gas-exchange calculations; therefore, the algorithm is expected to be useful in various models where the *A*_*n*_–*E*–*g*_*s*_ model is implemented, including vegetation, crop, ecosystem, hydrology, land-surface, climate, and Earth-system models (Sellers et al., 1992; Longo et al., 2019; Lawrence et al., 2019; Danabasoglu et al., 2020; Masutomi et al., 2016a; Nitta et al., 2020).

A second contribution is that the convergence rate of the proposed algorithm was quantified. In the numerical experiments, approximately 10 iterations were sufficient to obtain about 10^−3^ *μ*mol CO_2_ m^−2^ s^−1^ precision in *A*_*n*_, and approximately 20 iterations were sufficient to obtain about 10^−7^ *μ*mol CO_2_ m^−2^ s^−1^ precision. These levels corresponded to 0.1 ppm and 10^−5^ ppm precision in *C*_*i*_, respectively. To our knowledge, previous studies have not systematically evaluated convergence rates, so it is not yet possible to state whether the present method is faster or slower than other algorithms. Nevertheless, an iteration count on the order of 20 is computationally very small on current computers. The threshold analysis also showed that, up to the 10^−5^ ppm criterion tested here, the number of iterations increased approximately linearly with the number of decimal orders of precision required. Thus, the algorithm is not only guaranteed to converge, but also sufficiently fast for obtaining practical numerical precision in both *A*_*n*_ and *C*_*i*_.

A third implication is that the *F*–*G* formulation may be highly extensible. The present proof was developed for the Medlyn-type stomatal conductance model coupled with Farquhar-type C3 photosynthesis and explicit CO_2_ and H_2_O diffusion equations (Farquhar et al., 1980; Medlyn et al., 2011; Campbell and Norman, 1998). However, many variants of stomatal conductance models may be represented through different forms of *G*(*C*_*i*_) (Ball et al., 1987; Leuning, 1995; Damour et al., 2010), while the CO_2_ diffusion component that defines *F* (*C*_*i*_) often retains a similar mass-balance structure. Therefore, the same strategy may be applicable to other coupled gas-exchange models: derive the variant-specific *G*(*C*_*i*_), identify the appropriate solution interval, prove the ordering and monotonicity properties relative to *F* (*C*_*i*_), and then construct an iteration that necessarily approaches the unique crossing. This suggests that the present work is not only a solver for one specific formulation, but also a general framework for designing guaranteed-convergence algorithms for a wider class of leaf gas-exchange models.

Table 2 summarizes the conceptual differences between conventional iterative approaches and the proposed guaranteed-convergence algorithm, including the convergence guarantee and the convergence rate quantified in this study.

**Table 2:** Comparison of conventional iteration and the proposed algorithm.

| Feature | Conventional iteration | Proposed algorithm |
| --- | --- | --- |
| Iteration | Iterates over coupled internal variables | Iterates in one dimension, $C_i$ |
| Initial value | No general rule for choosing the initial value | Uses the lower endpoint of the analytically defined interval |
| Convergence | No general guarantee | Guaranteed |
| Convergence rate | Usually not systematically quantified | Rapid in the tested cases:<br>approximately 10 iterations for about $10^{-3} \mu\text{mol CO}_2 \text{ m}^{-2} \text{ s}^{-1}$ precision in $A_n$ and approximately 20 iterations for about $10^{-7} \mu\text{mol CO}_2 \text{ m}^{-2} \text{ s}^{-1}$ precision |

## 5 Conclusion

We developed a numerical algorithm whose convergence is guaranteed when solving the *A*_*n*_– *E*–*g*_*s*_ model. By eliminating concerns about nonconvergence, this algorithm provides robust and reliable leaf gas-exchange calculations for a wide range of models in which coupled photosynthesis, transpiration, and stomatal conductance are represented.

The convergence rate was also rapid in the numerical tests. Approximately 10 iterations were sufficient to obtain about 10^−3^ *μ*mol CO_2_ m^−2^ s^−1^ precision in *A*_*n*_, and approximately 20 iterations were sufficient to obtain about 10^−7^ *μ*mol CO_2_ m^−2^ s^−1^ precision. Together, guaranteed and rapid convergence make the algorithm practical across scales, from leaf-level gas-exchange calculations to large-scale simulations, including global-scale applications.

## A Temperature-response functions

Leaf temperature is denoted by *T*_*l*_ [K], and the reference temperature is *T*_0_ = 298.15 K. The Arrhenius factor is

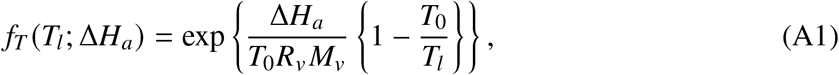

where *R*_*v*_ = 461.5 J kg^−1^ K^−1^ is the specific gas constant for water vapor, *M*_*v*_ = 18.01528 × 10^−3^ kg mol^−1^ is the molecular mass of water vapor, and Δ*H*_*a*_ [J mol^−1^] is the activation energy. The high-temperature inhibition factor is

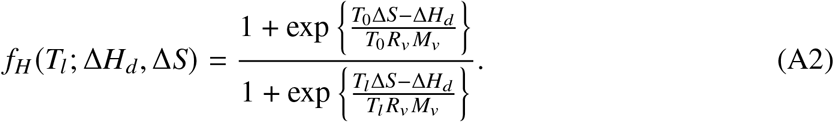

Here, Δ*H*_*d*_ [J mol^−1^] is the deactivation energy and Δ*S* [J mol^−1^ K^−1^] is the entropy term.

The Michaelis–Menten coefficients for CO_2_, *K*_*c*_ [Pa], and O_2_, *K*_*o*_ [Pa], and the CO_2_ compensation point, Γ^∗^ [ppm], are calculated as

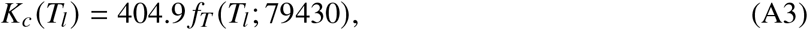

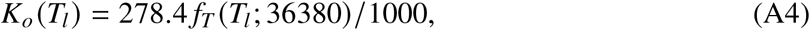

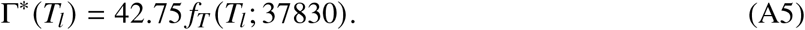

The maximum carboxylation rate, *V*_cmax_ [*μ*mol m^−2^ s^−1^], maximum electron transport rate, *J*_max_ [*μ*mol m^−2^ s^−1^], and day respiration, *R*_*L*_ [*μ*mol m^−2^ s^−1^], are

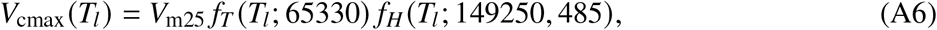

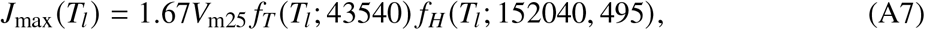

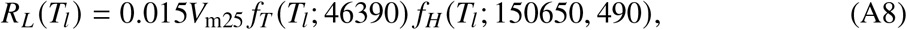

where *V*_m25_ [*μ*mol m^−2^ s^−1^] is the value of *V*_cmax_ at 25^◦^C. The electron transport rate used in the RuBP-regeneration-limited condition is obtained from the non-rectangular hyperbola

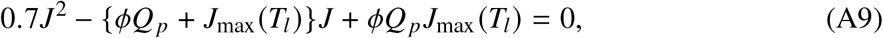

with the smaller positive root, where *ϕ* = 0.425 is the apparent quantum yield, representing the conversion efficiency from absorbed photosynthetic photon flux to electron transport, and *Q* _*p*_ [*μ*mol m^−2^ s^−1^] is the absorbed photosynthetic photon flux density.

In the numerical example, RH (relative humidity) = 80% is imposed by setting *e*_*a*_ = 0.8*e*_*i*_. Unless otherwise varied, the remaining parameter values are *V*_m25_ = 60 *μ*mol m^−2^ s^−1^, *g*_min_ = 0.01 mol m^−2^ s^−1^, and *g*_slp_ = 4.

## B Boundary-layer conductance *g*_*b*_

Following the aerodynamic resistance formulation of Allen et al. (1998), the boundary-layer conductance *g*_*b*_ is given by

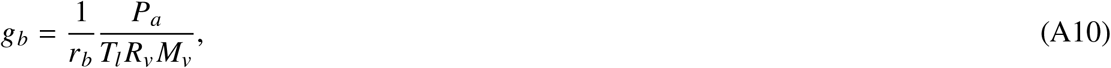

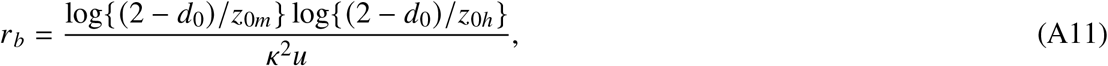

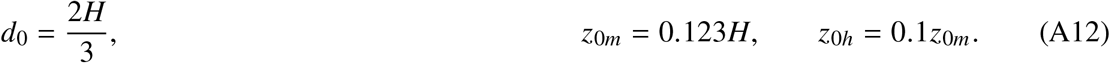

where *r*_*b*_ is the boundary-layer resistance, *P*_*a*_ is atmospheric pressure, *T*_*l*_ is leaf temperature, *R*_*v*_ = 461.5 J kg^−1^ K^−1^ is the specific gas constant for water vapor, *M*_*v*_ = 18.01528 × 10^−3^ kg mol^−1^ is the molecular mass of water vapor, *d*_0_ is the zero-plane displacement height, *z*_0*m*_ and *z*_0*h*_ are the roughness lengths for momentum and heat transfer, respectively, *κ* = 0.4 is the von Kármán constant, *u* is 2-m wind speed, and *H* is canopy height. In the numerical example, canopy height is set to *H* = 1 m unless otherwise varied.

## C Zeros and singularities for the function *F* (*C*_*i*_)

Under Rubisco- or RuBP-limited photosynthesis with *a* > 0, *F*_*R*_ (*C*_*i*_) = 0 is equivalent to *A*_*nR*_ (*C*_*i*_) = 0. Thus 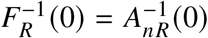, which is obtained from Eq. (11) as

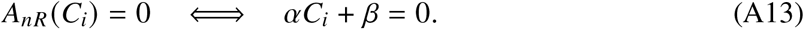

Therefore,

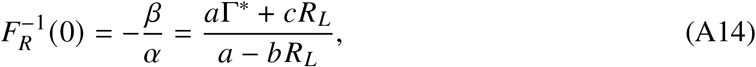

provided that *α* = *a* − *bR*_*L*_ ≠ 0. Equivalently, in terms of the original photosynthetic parameters,

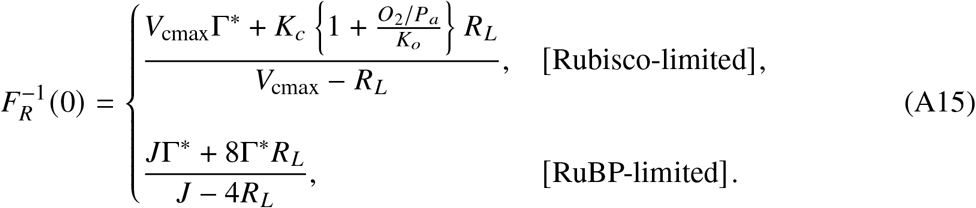

The singularities of *F* (*C*_*i*_) occur when its denominator is zero. For the Rubisco/RuBP-limited condition, substituting Eq. (11) into the denominator of Eq. (14) gives

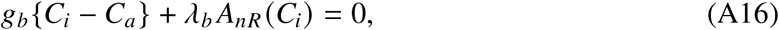

which is equivalent to the quadratic equation

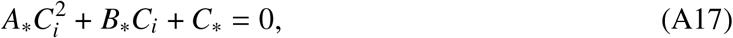

where

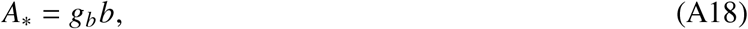

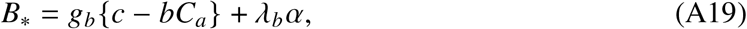

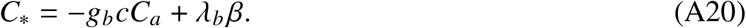

Thus, the two possible singular points are

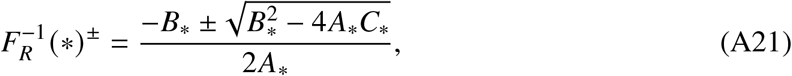

For the TPU-limited condition, the singular point follows directly from the denominator of Eq. (18):

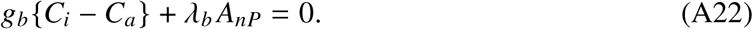

Thus,

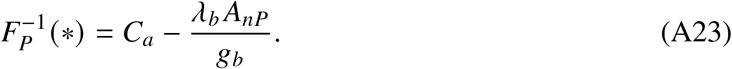

## D Inverse of the function *F* (*C*_*i*_)

For the Rubisco/RuBP-limited condition with *a* > 0, substituting Eq. (11) into *g*_*s*_ = *F*_*R*_ (*C*_*i*_) yields the quadratic equation

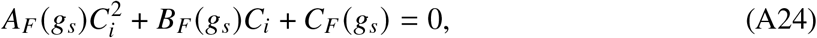

where

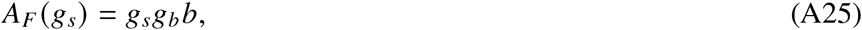

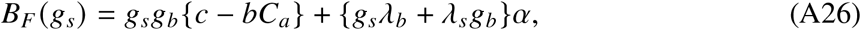

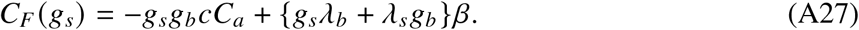

Hence the two possible inverse values are

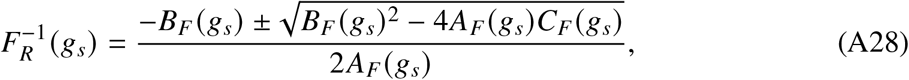

For the TPU-limited condition, *A*_*n*_ (*C*_*i*_) = *A*_*nP*_ = 3*T*_*p*_ − *R*_*L*_ is constant, and Eq. (18) can be inverted directly. Substituting *g*_*s*_ = *F*_*P*_ (*C*_*i*_) gives

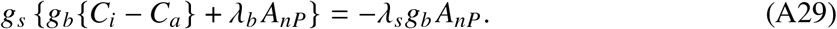

Thus, for *g*_*s*_ ≠ 0,

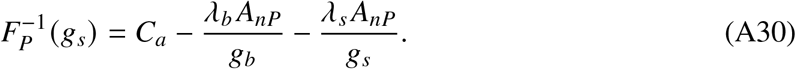

## E Iteration counts for the sensitivity tests

Figures A1–A5 summarize the number of iterations required for convergence for each value used in the one-variable sensitivity tests under convergence thresholds from 10^−5^ to 10^−1^ ppm, separately for the Rubisco-limited, RuBP-limited, and TPU-limited conditions. In each row, only the variable shown on the horizontal axis is varied; the three columns show the three limiting conditions, and all other environmental and physiological inputs are fixed at the Fig. 1 values.

**Figure A1:**
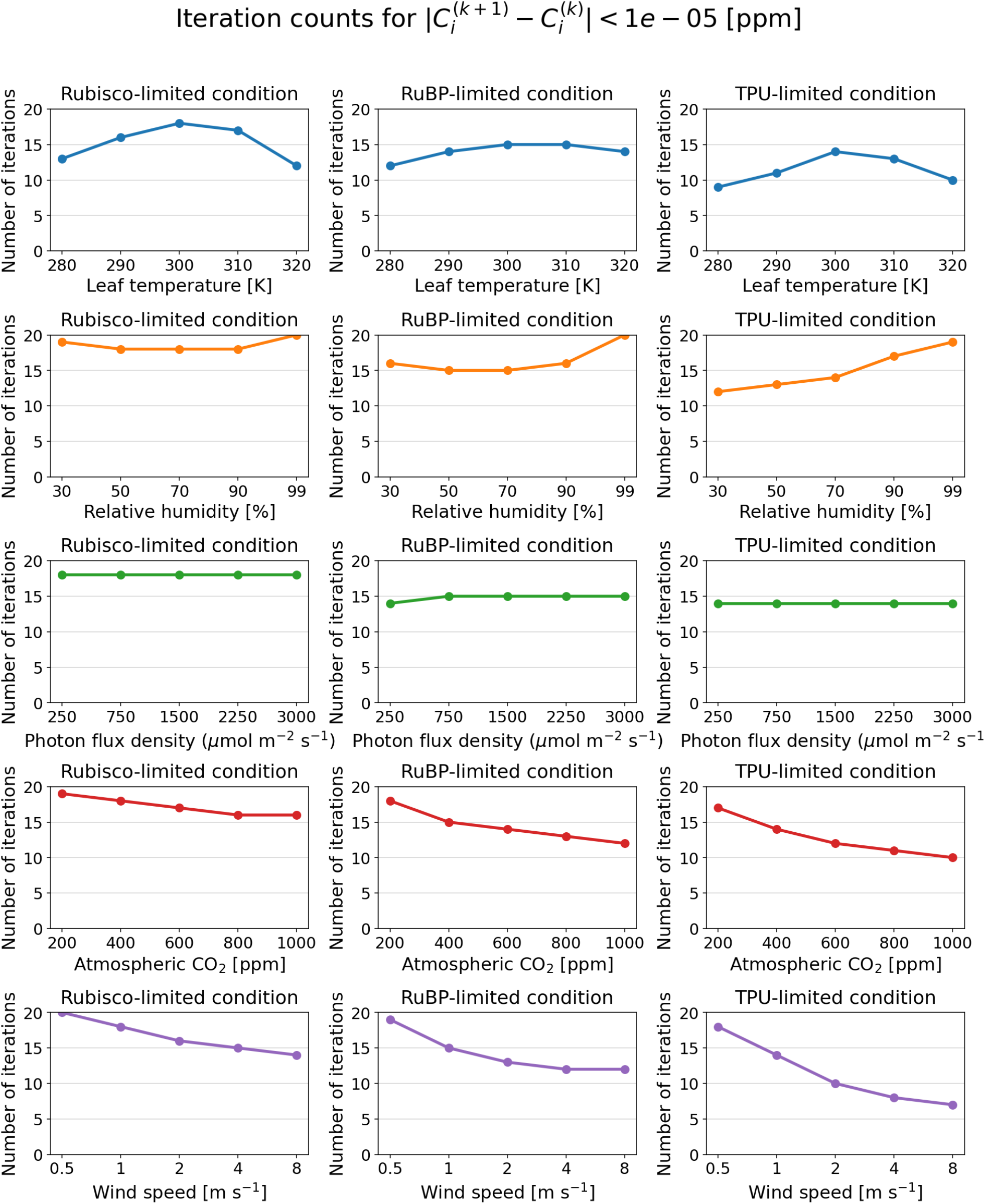
Number of iterations required for convergence in the one-variable sensitivity tests under the threshold 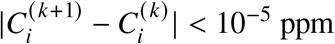.

**Figure A2:**
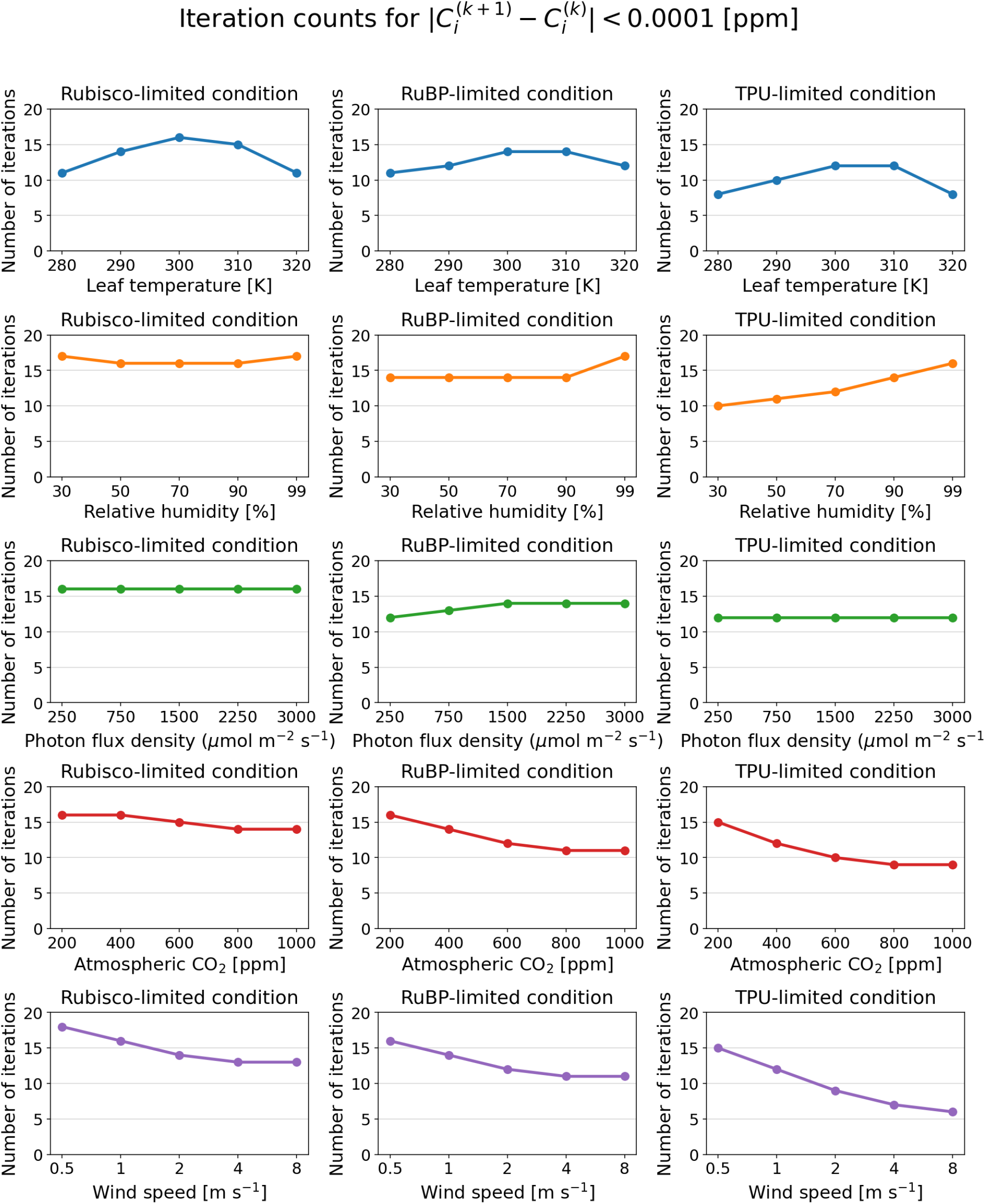
Number of iterations required for convergence in the one-variable sensitivity tests under the threshold 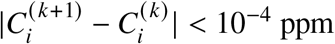.

**Figure A3:**
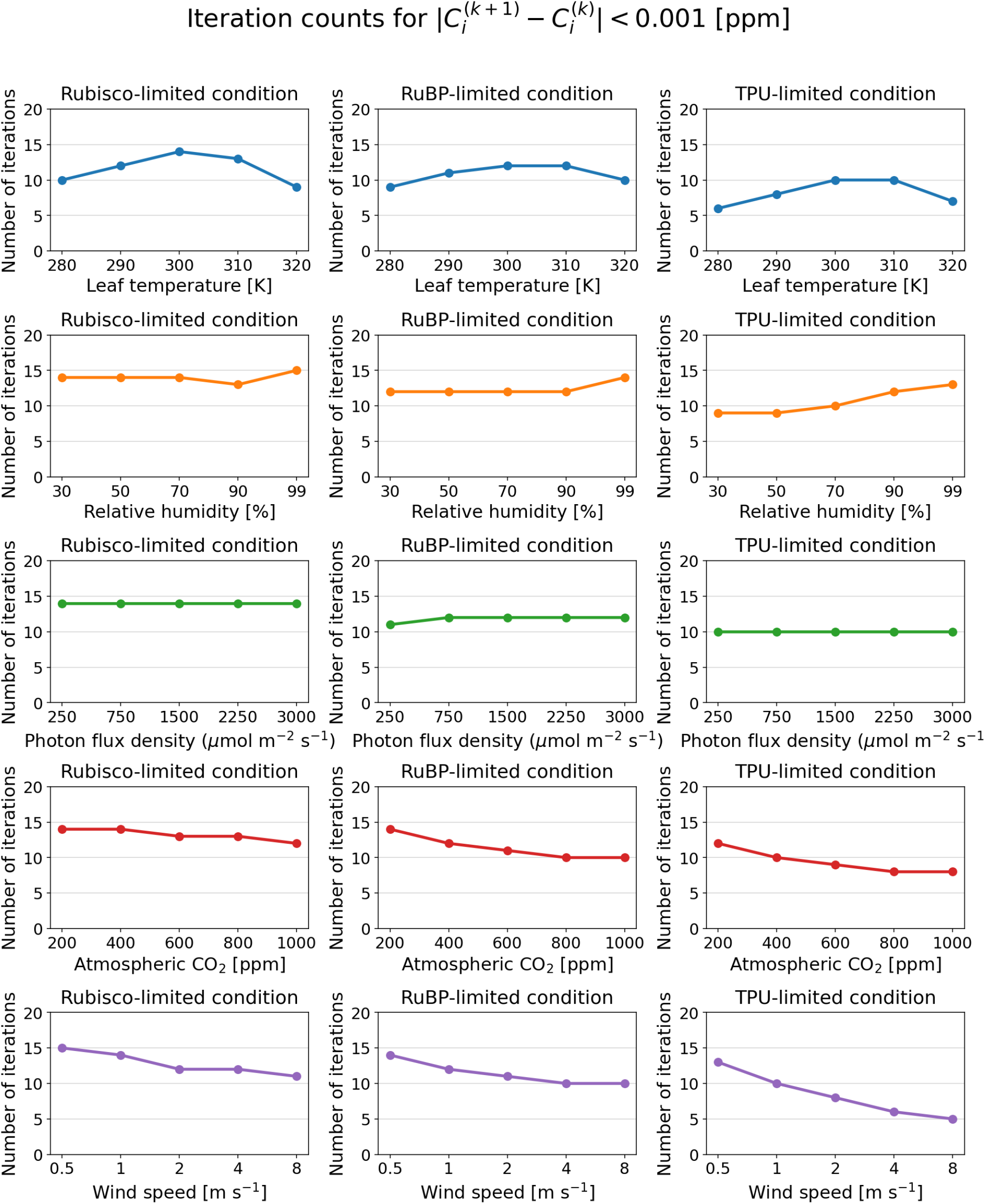
Number of iterations required for convergence in the one-variable sensitivity tests under the threshold 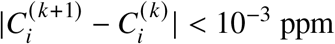.

**Figure A4:**
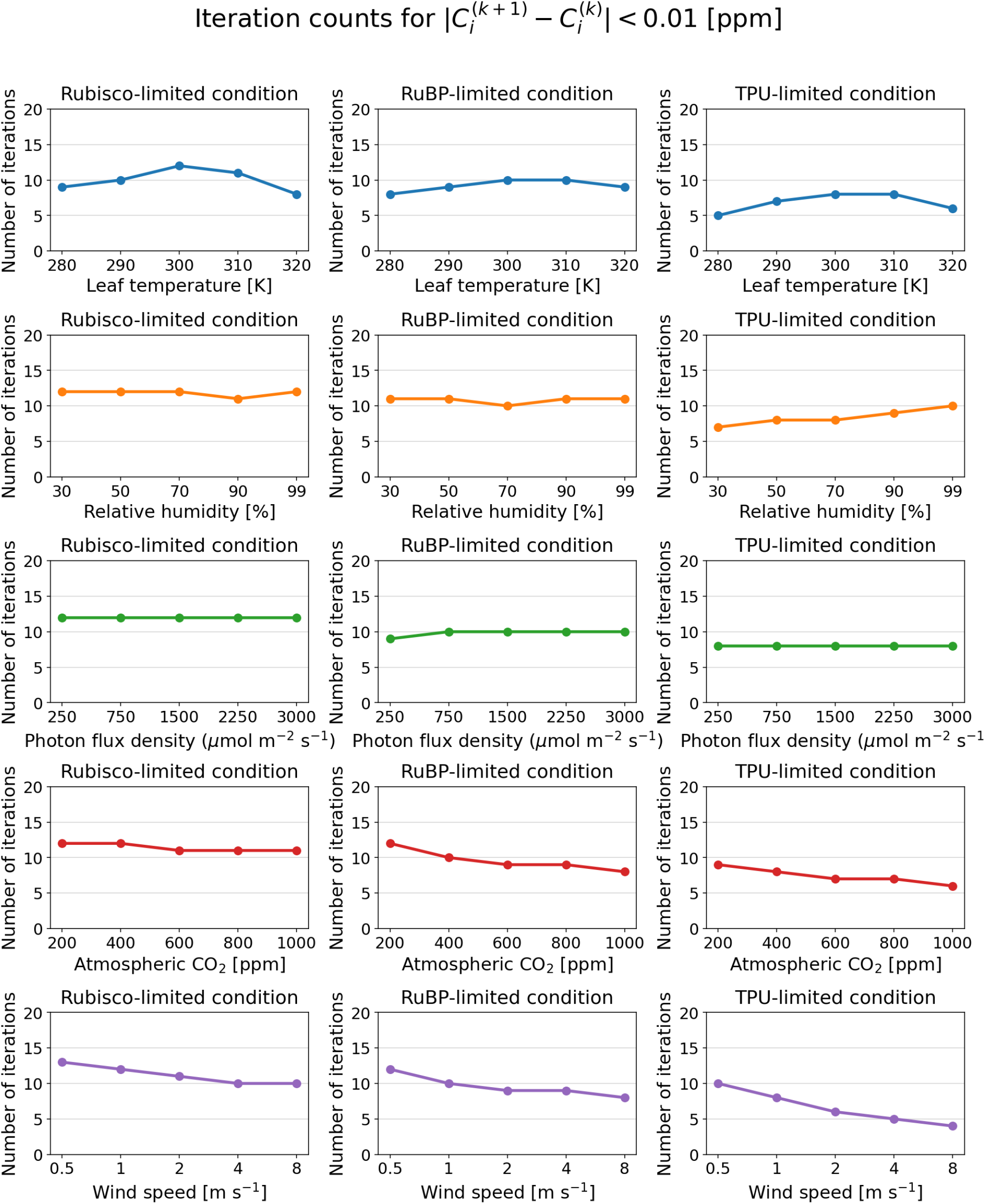
Number of iterations required for convergence in the one-variable sensitivity tests under the threshold 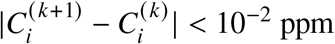.

**Figure A5:**
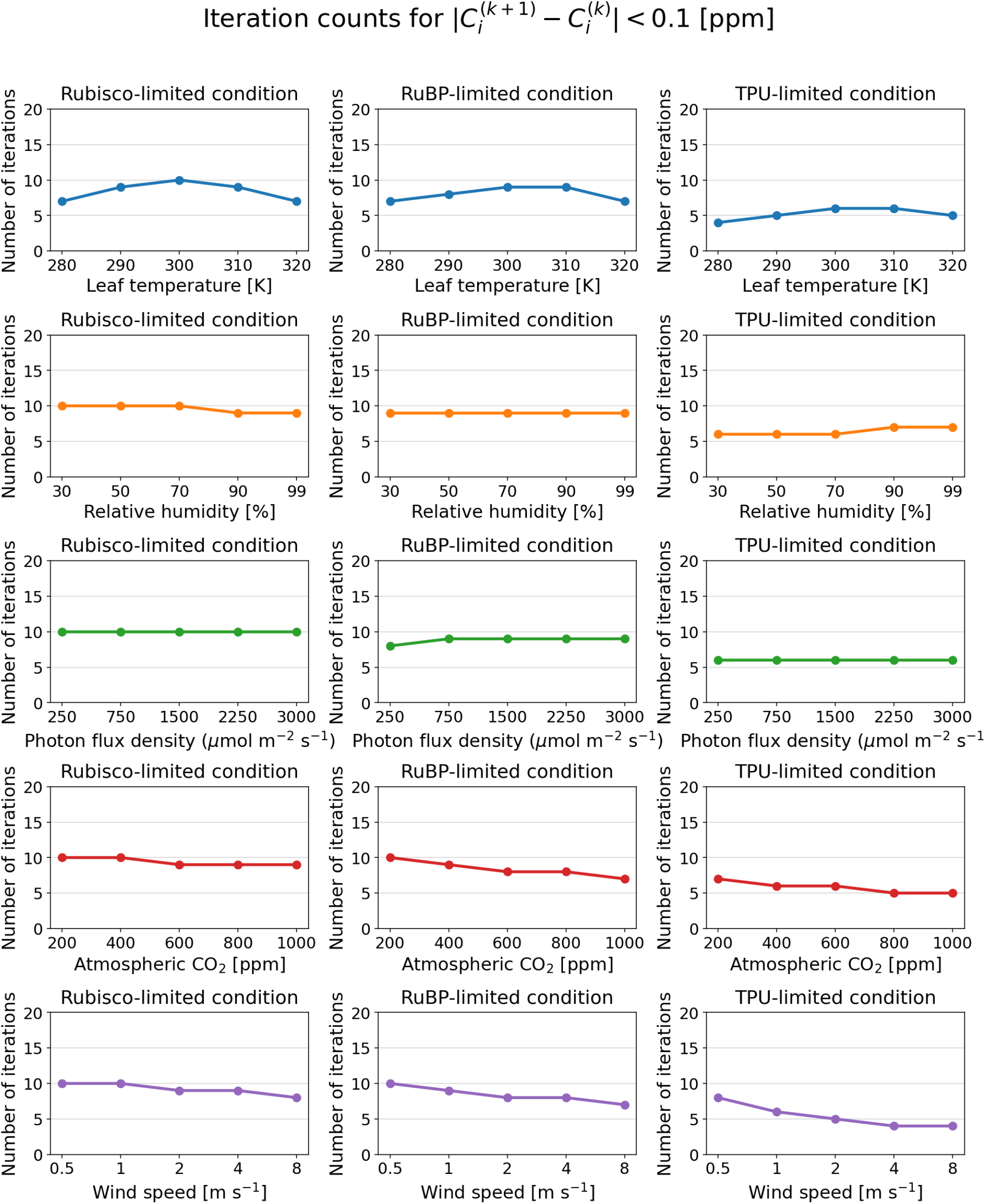
Number of iterations required for convergence in the one-variable sensitivity tests under the threshold 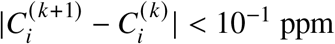.

## Acknowledgements

This study was partly supported by the Environment Research and Technology Development Fund (JPMEERF21S12020 and JPMEERF20252002) of the Environmental Restoration and Conservation Agency (ERCA), and JSPS KAKENHI Grant Number JP23H00351. We acknowledge the use of ChatGPT, developed by OpenAI, for assistance with translation from Japanese to English and with English-language editing. The authors reviewed and edited all AI-assisted text and take full responsibility for the content of the manuscript.

## Author Contributions

Y.M. conceived and designed the research, developed the algorithm, performed the analysis, interpreted the results, and drafted the manuscript. K.K. supervised the research and contributed to interpretation and manuscript revision.

## Data Availability

The equations and parameter values required to reproduce the model calculations are provided in the manuscript. The code and data supporting the numerical results will be archived in a public repository with a persistent identifier upon acceptance.

